# cGAS inhibition delays TDP-43-driven ALS Pathogenesis

**DOI:** 10.64898/2026.02.24.707791

**Authors:** Yajing Liu, Weixi Feng, Abulimiti Aikedan, Se-In Lee, Maitreyee Bhagwat, Ravi Kumar Nagiri, Man Ying Wong, Sadaf Amin, Wenhui Qu, Jingjie Zhu, Si-Yu Wang, Pearly Ye, Kendra Norman, Guillermo Coronas-Samano, Marta Olah, Hagen U Tilgner, Li Fan, Subhash C. Sinha, Li Gan

## Abstract

Amyotrophic lateral sclerosis (ALS) is a fatal neurodegenerative disorder marked by motor neuron loss and cytoplasmic mislocalization of TAR DNA-binding protein 43 (TDP-43), a key regulator of RNA splicing. However, the upstream modulators of this process remain poorly defined. Here we identify cyclic GMP–AMP synthase (cGAS) as a central mediator of TDP-43 pathology and associated mis-splicing. cGAS expression was elevated in ALS patient brains and enriched across activated microglia. In human iPSC-derived microglia–motor neuron co-cultures, neuronal TDP-43 pathology triggered microglial cGAS activation, whereas pharmacological inhibition with a potent human cGAS inhibitor reduced phosphorylated TDP-43, restored lysosomal and phagocytic programs, normalized microglial reactivity, and reversed TDP-43–associated RNA splicing defects. In vivo, cGAS inhibition in TDP-43 Q331K mice reversed widespread RNA splicing abnormalities across neurons and oligodendrocyte lineage cells, attenuated neurodegenerative pathology, and preserved motor function. Together, these findings identify cGAS as a druggable upstream regulator linking innate immune signaling to TDP-43–dependent RNA mis-splicing and neurodegeneration, and establish cGAS inhibition as a promising therapeutic strategy for ALS.

## Introduction

ALS is the most common adult-onset motor neuron disease, characterized by progressive degeneration of upper and lower motor neurons and a median survival of only two to four years after symptom onset^1–3^. The pathological hallmark of ALS is the cytoplasmic mislocalization and aggregation of TAR DNA-binding protein 43 (TDP-43), which occurs in more than 95% of patients^4,5^. TDP-43 pathology manifests as phosphorylated, ubiquitinated inclusions that disrupt multiple essential cellular processes, including RNA splicing, nucleocytoplasmic transport, and mitochondrial homeostasis^4,6–9^. Among these, TDP-43 loss of nuclear function or ALS-linked TDP-43 mutations could cause aberrant inclusion of cryptic exons in neuronal transcripts, directly contributing to neurodegeneration^8,10–14^. Restoring splicing fidelity has therefore emerged as a key therapeutic objective in TDP-43 proteinopathies.

Recent studies have implicated the cyclic GMP–AMP synthase (cGAS)–stimulator of interferon genes (STING) pathway in several neurodegenerative conditions, including ALS^15–18^. cGAS functions as a cytosolic DNA sensor that detects double-stranded DNA and catalyzes production of cyclic GMP–AMP (cGAMP), which activates STING to induce type I interferon signaling and inflammatory responses^19–21^. In ALS, pathological TDP-43 promotes mitochondrial injury and the release of mitochondrial DNA into the cytosol, thereby engaging the cGAS–STING pathway^16,22^. While STING inhibition has shown therapeutic benefit in ALS mouse models by reducing neuroinflammation and preserving motor function^16,18,23^, the upstream role of cGAS, the initiating and rate-limiting enzyme of this cascade, remains poorly understood.

Although cGAS and STING are canonically linked, accumulating evidence reveals that they mediate distinct biological processes beyond the interferon pathway. Nuclear cGAS regulates DNA repair and chromatin stability independently of STING by binding chromatin and suppressing homologous recombination^24,25^, whereas STING can be activated through interferon-γ–inducible factor 16 (IFI16) and ataxia telangiectasia-mutated protein (ATM) to drive NF-κB signaling or modulate cell-cycle progression independently of cGAS^26,27^. These findings highlight that selective inhibition of cGAS may achieve more targeted modulation of disease-relevant pathways than global suppression of STING signaling.

In the central nervous system (CNS), cGAS expression is largely restricted to microglia, positioning it as a key innate immune sensor linking neuroinflammation to neuronal dysfunction^28–30^. Consistent with this role, pharmacologic or genetic cGAS inhibition has shown neuroprotective effects in diverse contexts, including traumatic brain injury and Alzheimer’s disease^15,31,32^. cGAS inhibition reduces microglial activation, preserves synaptic integrity, and prevents cognitive decline^15^. Despite these promising findings, translation toward human disease has been hindered by two major challenges: the lack of potent inhibitors selective for human cGAS, and the absence of validated human cellular systems for mechanistic and therapeutic testing.

To address these gaps, we developed a potent, brain-permeable small molecule, SS-1386, that selectively inhibits human cGAS with nanomolar potency. Using human iPSC-derived microglia–motor neuron co-cultures, we show that neuronal TDP-43 pathology triggers microglial cGAS activation, leading to lysosomal dysfunction, inflammatory activation, and disruption of microglial phagocytic programs. Pharmacological inhibition of cGAS restored lysosomal and phagocytic function, reduced pathological TDP-43 phosphorylation, normalized microglial reactivity, and partially reversed TDP-43–associated RNA splicing defects in motor neurons. In vivo, treatment of TDP-43 Q331K mice with a mouse-selective cGAS inhibitor ameliorated motor deficits, preserved motor neurons, attenuated axonal injury, and corrected widespread TDP-43–dependent splicing dysregulation across neurons and oligodendrocyte lineage cells, thereby restoring RNA splicing homeostasis. In general, these findings identify cGAS as an upstream integrator of innate immune activation, glial dysfunction, and RNA dysmetabolism in ALS, and establish cGAS inhibition as a mechanistically grounded therapeutic strategy for TDP-43 proteinopathies.

## Results

### Microglial cGAS activation in human ALS tissues and iPSC-derived cellular model

To investigate cGAS involvement in ALS pathogenesis, we analyzed single-cell RNA-sequencing data from postmortem human ALS and non-ALS brains^33^. Initial unbiased clustering of all cell types identified seven major cell populations, including immune cells (clusters 1-6) and transcriptionally mixed cells (cluster 7) (**Extended Data Fig. 1a-c**). Given that cGAS expression in the CNS is predominantly microglial, we extracted immune clusters for high-resolution analysis and reclustered them into 11 distinct subclusters (**Fig. 1a**). Expression profiling of *CX3CR1* (C-X3-C motif chemokine receptor 1) and *CD3E* (CD3 epsilon subunit of T-cell receptor complex) identified subclusters 1-10 as microglia and subcluster 11 as T cells (**Extended Data Fig. 1d**).

**Figure 1.**
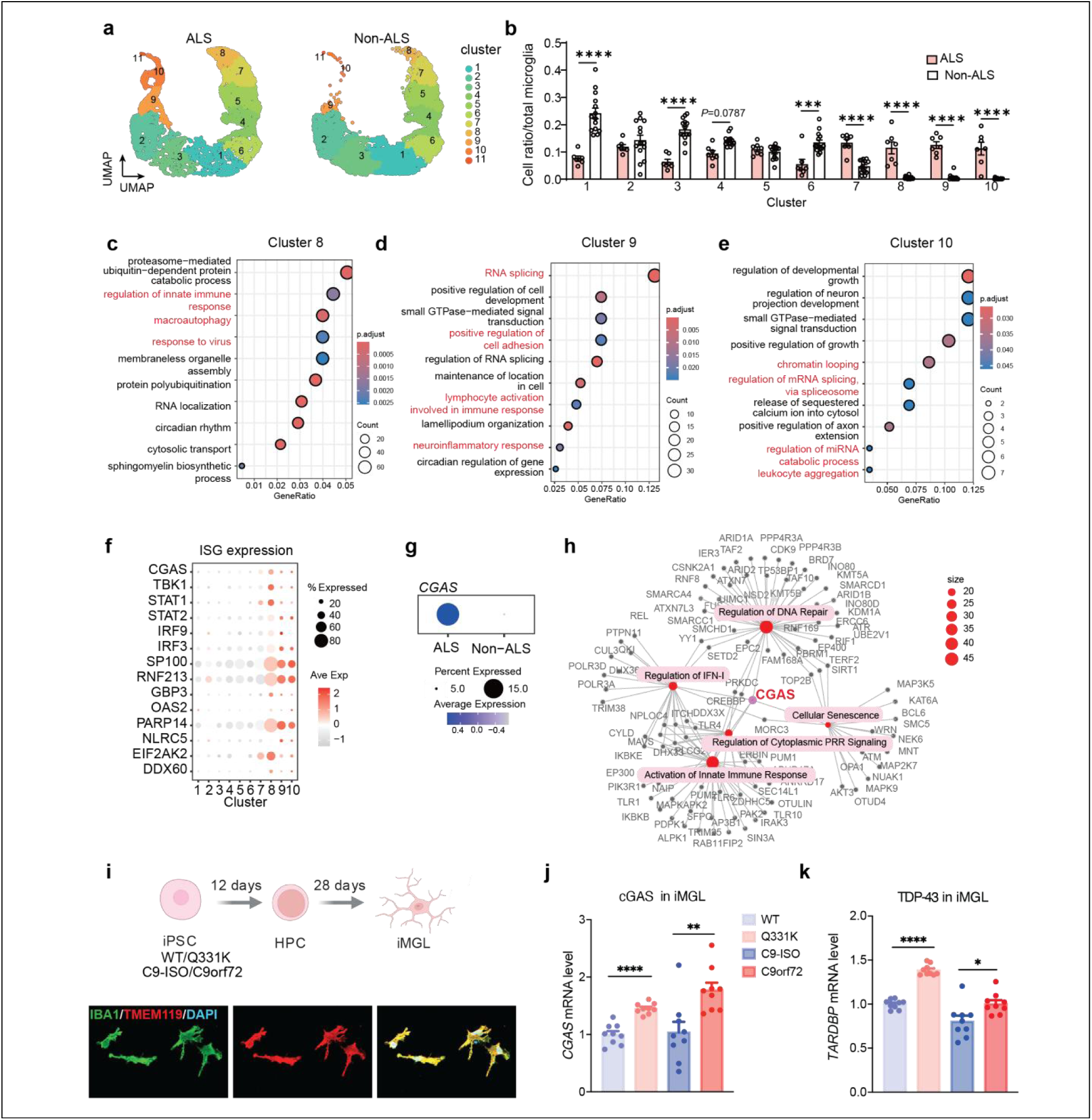
cGAS expression in microglia is elevated in human ALS brains and iPSC-derived microglia. **a,** Uniform Manifold Approximation and Projection (UMAP) plot of immune cells from the ALS and non-ALS brain samples. **b,** The cell ratio of each microglial subclusters between ALS samples and non-ALS samples. N = 7 for ALS samples and n = 14 for non-ALS samples. Data are presented as mean ± SEM. Cluster 1: \*\*\*\**P* < 0.0001; Cluster 3: \*\*\*\**P* < 0.0001; Cluster 4: *P* = 0.0787; Cluster 6: \*\*\**P* = 0.0001; Cluster 7: \*\*\*\**P* < 0.0001; Cluster 8: \*\*\*\**P* < 0.0001; Cluster 9: \*\*\*\**P* < 0.0001; Cluster 10: \*\*\*\**P* < 0.0001. Data were analyzed by two-way ANOVA with a Tukey multiple comparisons test. **c–e,** GO analysis of marker genes from microglial subcluster 8 (c), 9 (d), and 10 (e). **f,** Dotplot for the expression of ISGs in the microglial subclusters. **g,** Dotplot of *CGAS* expression in ALS microglia and non-ALS microglia. **h,** GO analysis for cGAS-associated pathways enriched in upregulated genes in ALS brain microglia. **i,** Diagram for the differentiation of iPSC-derived human microglial-like cells (iMGL) and representative immunofluorescence for the markers of iMGL (IBA1/TMEM119). **j,** *CGAS* mRNA levels of iMGLs in iPSC lines with ALS-mutations (WT, Q331K, C9- ISO, C9orf72). N = 9 for WT samples, n = 9 for Q331K samples, n = 9 for C9- ISO, and n = 9 for C9orf72 samples. Data are presented as mean ± SEM. Q331K vs. WT: \*\*\*\**P* < 0.0001; C9-ISO vs. C9orf72: \*\**P* = 0.0043. Data were analyzed by two-tailed unpaired t-test. **k,** *TARDBP* (TDP-43) mRNA levels of iMGLs in iPSC lines with ALS-mutations (WT, Q331K, C9- ISO, C9orf72). N = 9 for WT samples, n = 9 for Q331K samples, n = 9 for C9- ISO, and n = 9 for C9orf72 samples. Data are presented as mean ± SEM. Q331K vs. WT: \*\*\*\**P* < 0.0001; C9- ISO vs. C9orf72: \**P* = 0.0163. Data were analyzed by two-tailed unpaired t-test.

Analysis of microglial subcluster distribution revealed pronounced shifts in ALS brains, with microglia transitioning from homeostatic subclusters (1, 3, 4, and 6) toward disease-associated activated subclusters (7, 8, 9, and 10) (**Fig. 1b**). Gene ontology (GO) enrichment analysis of subcluster-specific marker genes revealed functional specialization, with subcluster 8 enriched for autophagy regulation and subclusters 9-10 enriched for RNA splicing pathways (**Fig. 1c-e**).

Notably, all ALS-enriched microglial subclusters exhibited significant enrichment of inflammatory response pathways, with subcluster 8 enriched for antiviral response pathways (**Fig. 1c-e**). The expression of interferon-stimulated genes (ISGs), a canonical hallmark of antiviral signaling, such as *TBK1*, *STAT1*, *STAT2*, *IRF3*, and *IRF9*, was markedly upregulated in subclusters 7-10, with particularly robust expression in subcluster 8 (**Fig. 1f)**, consistent with activation of interferon signaling cascades in ALS-associated microglia.

Given the central role of cGAS in initiating interferon responses, we next examined *CGAS* expression across microglial subclusters. *CGAS* was markedly elevated in ALS-enriched subclusters (7-10) relative to homeostatic subclusters, with highest expression in subcluster 8 (**Fig. 1f**), and showed significant upregulation in ALS microglia compared with non-ALS controls in a pseudobulk analysis (**Fig. 1g**). GO enrichment analysis further revealed that cGAS-associated pathways, including DNA repair regulation, type I interferon production, innate immune response activation, and cellular senescence, were significantly enriched among genes upregulated in ALS microglia (**Fig. 1h**). These findings implicate involvement of cGAS in driving aberrant microglial response in ALS.

To determine whether the interferon and cGAS signatures observed in human ALS microglia arise from cell-autonomous effects of disease-linked mutations, we turned to a genetically defined and tractable iPSC-based system. We generated iPSC-derived microglia-like cells (iMGLs) from lines carrying *TARDBP-Q331K* or *C9orf72* hexanucleotide expansions, two major genetic drivers of ALS that converge on TDP-43 pathology, together with matched isogenic controls. iMGLs expressed canonical microglial markers, including IBA1 and TMEM119 (**Fig. 1i**), confirming microglial identity. Consistent with human ALS tissue, *CGAS* mRNA was significantly elevated in ALS-mutant iMGLs relative to controls (**Fig. 1j**), accompanied by increased *TARDBP* expression (**Fig. 1k**). These findings demonstrate that ALS-associated mutations can activate microglial cGAS signaling in a cell-autonomous manner.

### cGAS inhibition restores lysosomal function and enhances microglial clearance

Given the prominent activation of cGAS interferon signaling in ALS-associated microglia, we next asked whether pharmacological inhibition of cGAS could reverse disease-relevant microglial dysfunction. To this end, we developed SS-1386, a potent human-selective cGAS inhibitor with an IC50 of 0.071 μM (**Fig. 2a, Extended Data Fig. 2a**). The specificity of SS-1386 was confirmed using *CGAS*^-/-^ line, in which HT-DNA–induced CXCL10 response was abolished, and no additional effect of the inhibitor was observed (**Fig. 2b**). Pharmacological inhibition of cGAS with SS-1386 markedly attenuated downstream TBK1 activation, as evidenced by reduced TBK1 phosphorylation in immunoblotting (**Fig. 2c, d**), and suppressed IFN responses (Extended Data **Fig. 2b**).

**Figure 2.**
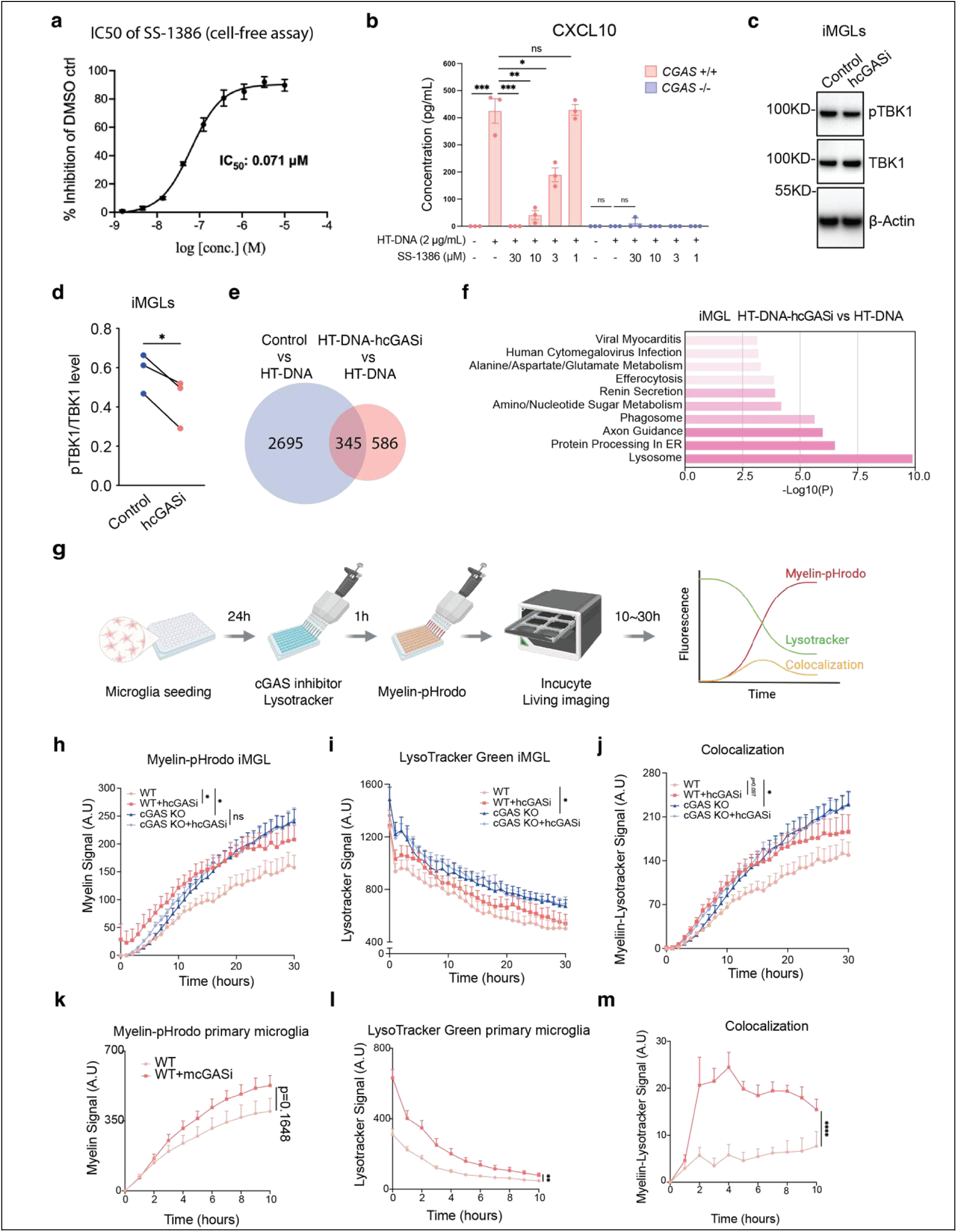
Human cGAS inhibitor improves phagocytosis and lysosomal mass in human iMGLs. **a,** Dose-dependent inhibition of SS-1386 on human cGAS/HT-DNA catalyzed conversion of ATP and GTP to produce cGAMP evaluated by quantifying the remaining ATP concentrations in the reactions using ATP Glo assay. IC50 = 0.071 µM. Data are reported as mean ± SEM. N = 3 per condition**. b,** Quantification of CXCL10 protein by ELISA in culture medium supernatants of *CGAS* ^+/+^ and *CGAS*^−/−^ THP-1 cells treated with HT-DNA and SS-1386. N = 3 for *CGAS* ^+/+^ samples, and n = 3 for *CGAS* ^-/-^ samples. Data are presented as mean ± SEM. HT-DNA vs. Control: \*\*\**P* = 0.0007; HT-DNA-30μM SS-1386 vs. HT-DNA: \*\*\**P* = 0.0007; HT-DNA-10μM SS-1386 vs. HT-DNA: \*\**P* = 0.0013; HT-DNA-3μM SS-1386 vs. HT-DNA: \**P* = 0.0105. Data were analyzed by two-way ANOVA with a Tukey multiple comparisons test; ns, not significant. **c and d**, Representative immunoblots (**c**) and quantification (**d**) of pTBK1 and TBK1 antibodies in iMGL with or without human cGAS inhibitor (hcGASi) normalized to β-Actin. N = 3 for Control samples, and n = 3 for hcGASi samples. hcGASi vs. Control: \**P* = 0.0322. Data were analyzed by two-tailed paired t-test. **e**, Venn diagram for gene sets in iMGLs comparing Control vs. HT-DNA and HT-DNA-hcGASi vs. HT-DNA. **f,** Pathway enrichment bar plot for iMGL transcriptomes of HT-DNA-hcGASi vs. HT-DNA. **g**, Experimental schematic of Incucyte live imaging for iMGL with myelin-pHrodo Red and LysoTracker. **h–j.** Time-course analysis of cGAS inhibition or genetic deletion increased myelin–pHrodo signal (h), LysoTracker Green signal (i), and myelin–lysosome colocalization (j) in iMGLs (n = 10 per group). Statistical analysis by two-way ANOVA with Tukey multiple comparisons; ns, not significant. Data are presented as mean ± SEM. myelin-pHrodo signal, WT + hcGASi vs. WT: \**P* = 0.0266; cGAS KO vs. WT: \**P* = 0.0307; LysoTracker Green signal, cGAS KO vs. WT: \**P* = 0.0431; cGAS KO vs. WT: \**P* = 0.0275. **k–m,** Time-course analysis of primary microglia (WT, WT+mcGASi). N = 6 for WT samples, and n = 6 for WT + mcGASi samples. Data are presented as mean ± SEM. k, myelin-pHrodo signal, WT + mcGASi vs. WT: *P* = 0.1648; l, LysoTracker Green signal, WT + mcGASi vs. WT: \*\**P* = 0.0017; m, myelin–lysosome colocalization, WT + mcGASi vs. WT: \*\*\*\**P* < 0.0001. Data were analyzed by two-way ANOVA with a Tukey multiple comparisons test.

We next assessed how cGAS inhibition altered microglial transcriptional and functional programs. Bulk RNA sequencing of iMGLs under control, HT-DNA, and HT-DNA plus hcGASi conditions revealed substantial overlap between genes induced by HT DNA and those reversed by hcGASi treatment (**Fig. 2e**). Gene set enrichment analysis showed that hcGASi reversed HT DNA–driven lysosomal and phagosomal pathways (**Fig. 2f**), implicating cGAS as a central regulator of microglial degradative function. To functionally validate these changes, we performed live cell imaging using LysoTracker Green and pHrodo Red–conjugated myelin debris to quantify lysosomal activity, phagocytosis, and acidification in real time (**Fig. 2g**). Over 30 h, hcGASi treated and cGAS knockout iMGLs exhibited increased pHrodo, LysoTracker, and colocalization signals relative to controls (**Fig. 2h–j**), indicating enhanced clearance upon cGAS inhibition. Similar effects were observed in primary mouse microglia treated with the mouse cGAS inhibitor TDI-6570, a mouse cGAS inhibitor reported previously^15^ (**Fig. 2k–m**), demonstrating conserved cGAS dependent regulation of microglial clearance across species.

### cGAS inhibition rescued functional and metabolic dysfunction in ALS mouse models

Building on our identification of robust cGAS activation in microglia from human ALS brains and its recapitulation in ALS mutant human iPSC-derived microglia-like cells, we next asked whether pharmacological inhibition of cGAS could provide therapeutic benefit in vivo. We tested this hypothesis in TDP-43 Q331K transgenic mice, a well-established ALS model that exhibits progressive motor neuron degeneration and TDP-43 pathology^10^. Because the human-selective inhibitor SS-1386 does not inhibit murine cGAS, we employed the validated mouse cGAS inhibitor TDI-6570^15,34^. Mice were placed on a TDI-6570–containing diet from 6 weeks of age, prior to symptom onset, through 32 weeks (**Fig. 3a**). Behavioral and physiological outcomes were assessed in WT, Q331K-control, and Q331K-cGASi cohorts (**Fig. 3a**). Q331K-control mice exhibited progressive motor impairment in accelerating rotarod assays performed between 20 and 28 weeks, whereas cGAS inhibition significantly preserved motor coordination and endurance (**Fig. 3b**).

**Figure 3.**
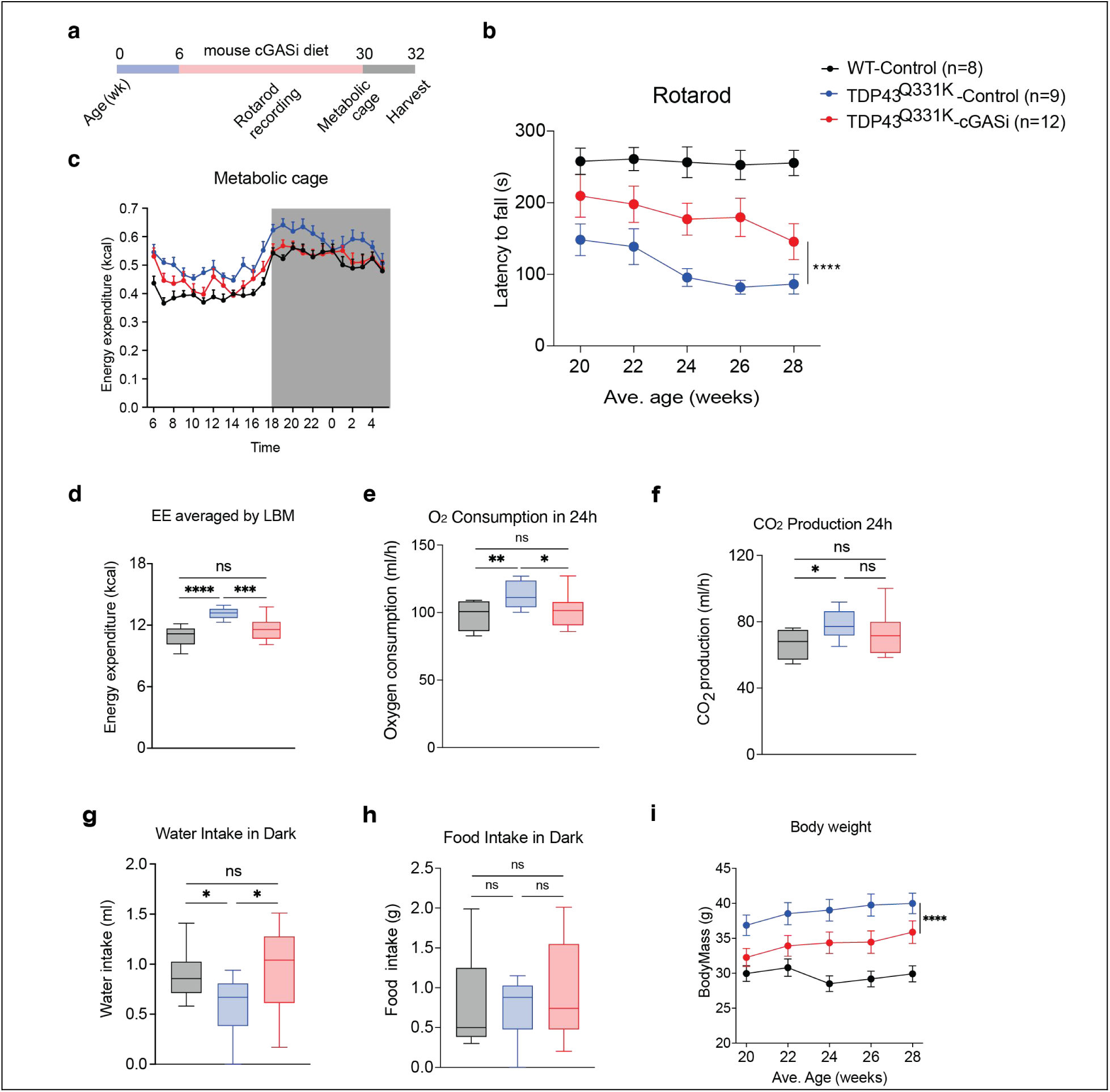
Mouse cGAS inhibitor alleviates motor and metabolic defect in Q331K ALS mouse model. **a,** Experimental schematic for mouse cGAS inhibitor diet, behavior recording, and tissue harvest for Q331K mice. **b,** Rotarod latency to fall (s) across ages from 20 weeks to 28 weeks for WT, Q331K-Control, and Q331K-cGASi groups. N = 8 for WT-Control samples, n = 9 for Q331K-Control samples, and n = 12 for Q331K-cGASi samples. Data are presented as mean ± SEM. Q331K-cGASi vs. Q331K-Control: \*\*\*\**P* < 0.0001. Data were analyzed by two-way ANOVA with a Tukey multiple comparisons test. **c,** Energy expenditure (EE) (kcal) recorded in metabolic cages over a continuous 24-h period for WT, Q331K-Control, and Q331K-cGASi. N = 8 for WT-Control samples, n = 9 for Q331K-Control samples, and n = 12 for Q331K-cGASi samples. Data are presented as mean ± SEM, and were analyzed by one-way ANOVA followed by a Tukey multiple comparison test (d-h); ns, not significant. **d,** EE normalized to lean body mass (LBM) (kcal) of WT, Q331K-Control, and Q331K-cGASi mice recorded in metabolic cages over 24 h. Q331K-Control vs. WT-Control: \*\*\*\**P* < 0.0001; Q331K-cGASi vs. Q331K-Control: \*\*\**P* = 0.0001. **e**, Oxygen consumption (ml/h) of WT, Q331K-Control, and Q331K-cGASi mice recorded in metabolic cages over 24 h. Q331K-Control vs. WT-Control: \*\**P* = 0.0081; Q331K-cGASi vs. Q331K-Control: \**P* = 0.0285. **f,** CO₂ production (ml/h) of WT, Q331K- Control, and Q331K-cGASi mice recorded in metabolic cages over 24 h. Q331K-Control vs. WT-Control: *P* = 0.0162. **g,** Water intake (ml) of WT, Q331K-Control, and Q331K-cGASi mice recorded in metabolic cages during the dark phase. Q331K-Control vs. WT-Control: \**P* = 0.0443; Q331K-cGASi vs. Q331K-Control: \**P* = 0.0360. **h,** Food intake (**g**) of WT, Q331K- Control, and Q331K-cGASi mice recorded in metabolic cages during the dark phase. **i**, Body weight of WT, Q331K- Control, and Q331K-cGASi mice across age points from 20 weeks to 28 weeks. N = 8 for WT-Control samples, n = 9 for Q331K-Control samples, and n = 12 for Q331K-cGASi samples. Data are presented as mean ± SEM. Q331K-cGASi vs. Q331K-Control: \*\*\*\**P* < 0.0001. Data were analyzed by two-way ANOVA with a Tukey multiple comparisons test.

In addition to motor deficits, TDP-43 Q331K mice are reported to develop systemic metabolic dysregulation, including obesity and elevated energy expenditure^35^. To assess systemic metabolic consequences of cGAS inhibition, we performed comprehensive metabolic profiling in Q331K mice using indirect calorimetry. Q331K-control mice exhibited increased total energy expenditure compared with WT mice over a 24 h period (**Fig. 3c**), which remained significant after normalization to lean body mass (**Fig. 3d**). cGAS inhibition significantly reduced this elevated energy expenditure, restoring values toward WT levels (**Fig. 3d**). Consistent with this normalization, Q331K-control mice showed increased oxygen consumption and CO₂ production relative to WT animals, both of which were significantly attenuated by cGAS inhibition (**Fig. 3e, f**). In parallel, Q331K mice displayed decreased water intake during the dark phase, which was normalized by cGASi treatment, while food intake remained unchanged across groups (**Fig. 3g, h**). Longitudinal monitoring further revealed excessive weight gain in Q331K mice between 20 and 28 weeks of age, which was markedly attenuated by cGAS inhibition (**Fig. 3i**). Together, these data indicate that cGAS inhibition restores metabolic efficiency and mitigates systemic metabolic dysregulation associated with TDP-43 pathology.

cGAS inhibition similarly delayed disease progression and reduced hindlimb paralysis in a second TDP-43–based ALS model (TARDBP-A315T)^36^ (**Extended Data Fig. 3a, b**), but had no protective effect in SOD1-G93A mice (**Extended Data Fig. 3c, d**)^37^. These results indicate that cGAS modulation selectively mitigates TDP-43–driven pathology rather than generalized motor neuron degeneration.

### cGAS inhibition ameliorates neuronal TDP-43 pathology and protects motor neurons in ALS mouse and human models

Having established that cGAS inhibition ameliorates TDP-43-mediated functional deficits, we next examined its impact on neuronal pathology in the ALS model. Immunoblot analysis of spinal cord lysates revealed that the treatment with mouse cGAS inhibitor significantly reduced phosphorylated TDP-43 (pTDP-43) levels in Q331K-cGASi mice compared to Q331K mice (**Fig. 4a, b**), indicating that cGAS signaling drives pathological TDP-43 modification in vivo. Beyond phospho-TDP-43, neurofilament light chain (NFL) serves as a sensitive biomarker of neuronal and axonal damage and is elevated in the vast majority of TDP-43 proteinopathy cases in ALS. ELISA quantification revealed that Q331K-Control mice displayed markedly elevated serum NFL concentrations relative to WT mice, with cGAS inhibition providing partial rescue (**Fig. 4c**).

**Figure 4.**
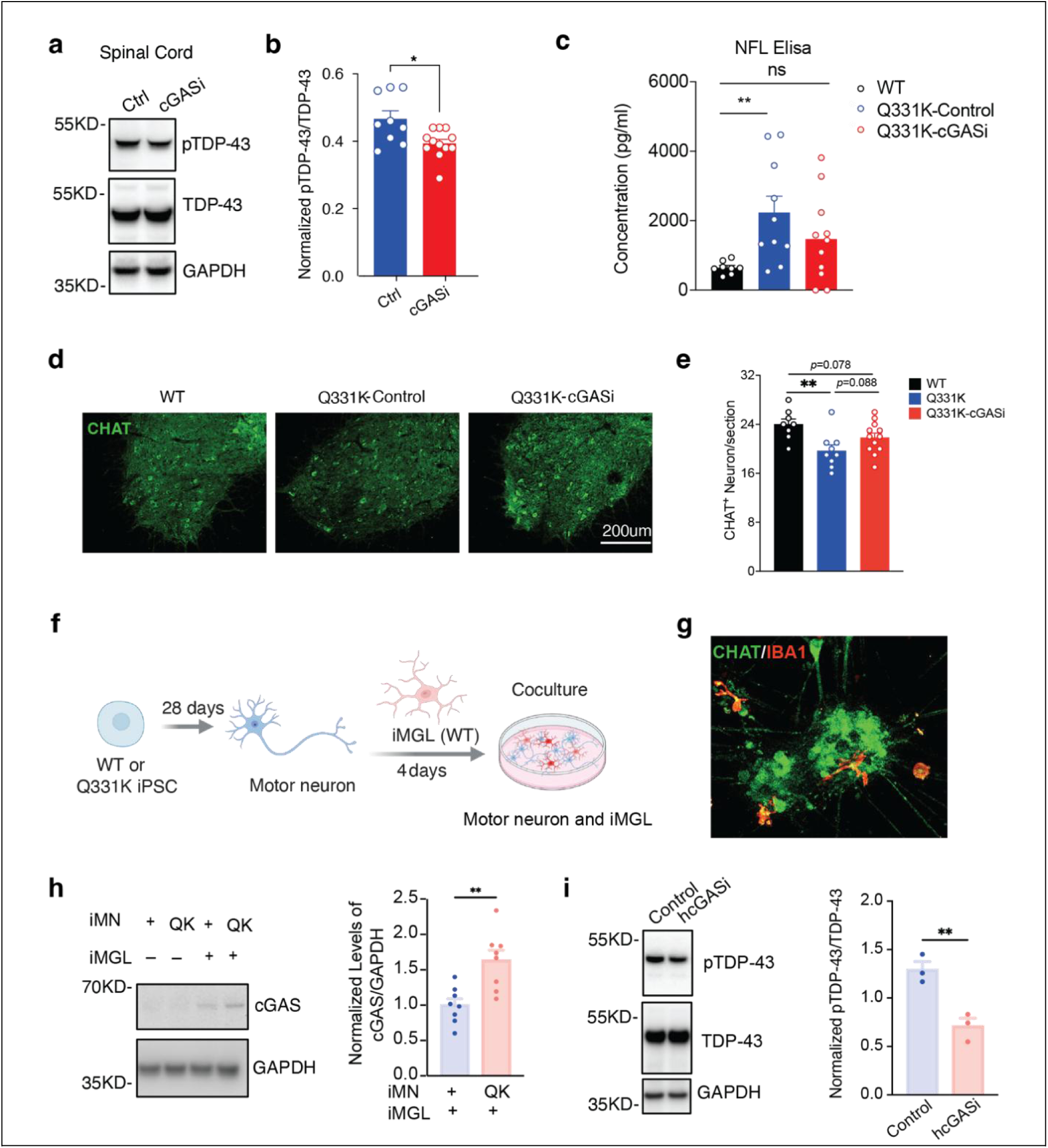
cGAS inhibitor alleviates TDP-43 pathology and the loss of motor neuron in mouse and human models of ALS. **a and b,** Representative immunoblots (a) and quantification (b) of pTDP-43 and TDP-43 antibodies in spinal cord of Q331K with or without cGASi normalized to GAPDH. N = 9 for Q331K-Control samples, and n = 12 for Q331K-cGASi samples. Data are presented as mean ± SEM. Q331K-cGASi vs. Q331K-Control: \**P* = 0.0105. Data were analyzed by two-tailed unpaired t-test. **c,** NFL ELISA for serum samples of WT, Q331K-Ctrl, and Q331K-cGASi mice. N = 8 for WT-Control samples, n = 10 for Q331K-Control samples, and n = 11 for Q331K-cGASi samples. Data are presented as mean ± SEM. Q331K-Control vs. WT-Control: \*\**P* = 0.0084. Data were analyzed by one-way ANOVA followed by a Tukey multiple comparison test; ns, not significant. **d and e,** Representative immunofluorescence images (d) and quantification (e) of CHAT⁺ neurons in spinal cord of WT, Q331K-Ctrl, and Q331K-cGASi groups. N = 8 for WT-Control samples, n = 9 for Q331K-Control samples, and n = 12 for Q331K-cGASi samples. Data are presented as mean ± SEM. Q331K-Control vs. WT-Control: \*\**P* = 0.0093; Q331K-cGASi vs. Q331K-Control: *P* = 0.0879; Q331K-cGASi vs. WT-Control: *P* = 0.0775. Data were analyzed by one-way ANOVA followed by a Tukey multiple comparison test. **f and g,** Diagram for co- culture of motor neurons and iMGLs and representative immunofluorescence for the markers of motor neuron (CHAT) and iMGL (IBA1). **h,** Representative immunoblots and quantification of cGAS antibody in the co- culture of motor neurons (WT and Q331K) and iMGLs normalized to GAPDH. N = 8 for co- culture-WT samples, and n = 8 for co- culture-Q331K samples. Data are presented as mean ± SEM. co- culture-Q331K vs. co culture-WT: \*\**P* = 0.0024. Data were analyzed by two-tailed unpaired t-test. **i,** Representative immunoblots and quantification of pTDP-43 and TDP- 43 antibodies in the co- culture of Q331K motor neurons and iMGLs with or without human cGAS inhibitor (hcGASi) normalized to GAPDH. N = 3 for Control samples, and n = 3 for hcGASi samples. Data are presented as mean ± SEM. hcGASi vs. Control: \*\**P* = 0.0078. Data were analyzed by two-tailed unpaired t-test.

To assess functional outcomes, we quantified motor neuron survival using CHAT immunofluorescence. Q331K-Control mice exhibited marked loss of CHAT^+^ neurons (19.7 neurons/section) compared to WT controls (24 neurons/section), whereas cGAS inhibition preserved motor neuron populations (21.8 neurons/section) (**Fig. 4d, e**). Together, these findings demonstrate that cGAS inhibition suppresses TDP-43 pathology, preserves motor neurons, and confers broad neuroprotection in ALS.

We next established the relevance of these findings in an all-human neuron–microglia context. We co-cultured wild-type iMGLs with iPSC-derived motor neurons expressing either *TARDBP* Q331K or wild-type *TARDBP*. Co-immunostaining confirmed distinct neuronal (CHAT⁺) and microglial (IBA1⁺) populations within the co-culture system (**Fig. 4f, g**). Immunoblot analysis revealed that Q331K motor neurons induced a marked upregulation of cGAS protein in co-cultured microglia (**Fig. 4h**), demonstrating that neuronal TDP-43 pathology directly activates microglial cGAS signaling. Consistent with the findings in animal model, SS-1386 treatment markedly reduced pTDP-43 levels in Q331K motor neuron–iMGL co-cultures (**Fig. 4i**), supporting that cGAS inhibition disrupts the reciprocal pathological signaling between diseased neurons and reactive microglia.

### cGAS inhibition reverses disease-associated microglial activation of ALS mice

To explore how cGAS inhibition disrupts reciprocal pathological signaling between neurons and microglia, we performed single-nucleus RNA sequencing (snRNA-seq) of spinal cords from WT, Q331K-control, and Q331K-cGASi mice (**Extended Data Fig. 4a-d**). Both the Q331K mutation and cGAS inhibition altered the relative abundance of major spinal cord cell types, including microglia, oligodendrocyte lineage cells, and neurons, prompting us to further characterize cell-type-specific transcriptional changes (**Extended Data Fig. 4e, f**). As cGAS is most highly expressed in microglia, we first examined the effects of Q331K and cGASi on microglial states. Q331K mice showed a marked expansion of the microglial population (8.2% of total nuclei vs. 4.1% in WT), which was partially normalized by mcGASi treatment (5.9%) (**Fig. 5a**), indicating that cGAS signaling promotes microglial proliferation or recruitment during disease progression.

**Figure 5.**
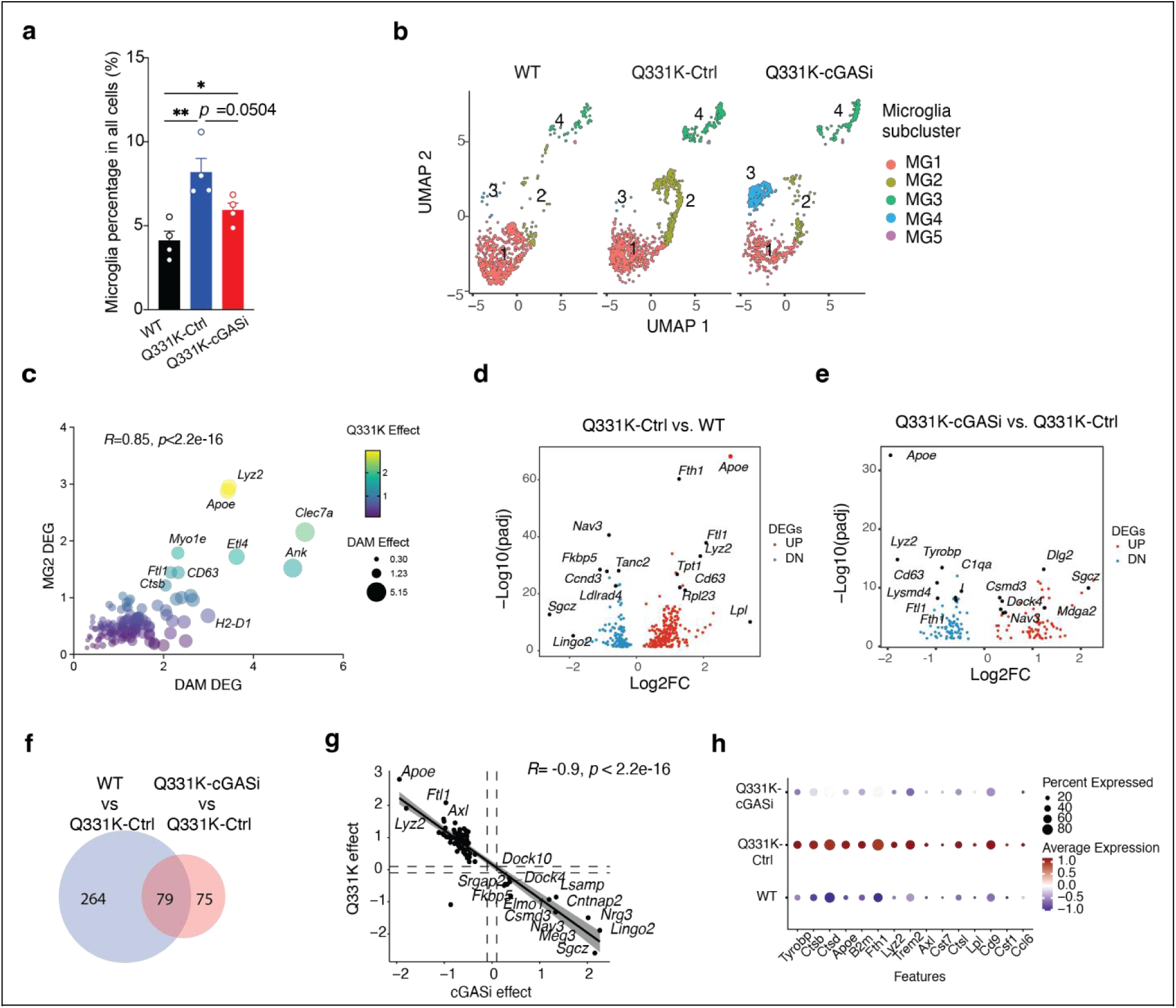
Mouse cGAS inhibitor restores microglial homeostasis in ALS mice. **a,** The percentage of microglia in the total cells of spinal cord in the single-nucleus analysis (WT, Q331K-Ctrl, Q331K-cGASi). N = 4 for WT-Control samples, n = 4 for Q331K-Control samples, and n = 4 for Q331K-cGASi samples. Data are presented as mean ± SEM. Q331K-Control vs. WT-Control: \*\**P* = 0.0042; Q331K-cGASi vs. Q331K-Control: *P* = 0.0504; Q331K-cGASi vs. WT-Control: \**P* = 0.0404. Data were analyzed by one-way ANOVA followed by a Tukey multiple comparison test. **b,** UMAP plot of microglial subclusters (MG1–MG5) across the three cohorts. **c,** Bubble plot for the levels of DAM-associated genes in MG2 subcluster. **d and e,** Volcano plot for microglial DEGs in the comparison of Q331K-Ctrl vs. WT (d) or Q331K-cGASi vs. Q331K-Ctrl (e). **f,** Venn diagram for microglial gene sets in the comparisons of WT vs. Q331K-Ctrl and Q331K-cGASi vs. Q331K-Ctrl. **g,** Correlation scatter plot with linear regression for microglial DEGs under cGASi effect and Q331K effect. **h,** Dotplot for ALS response associated marker genes in microglia of spinal cord in WT, Q331K-Ctrl, and Q331K-cGASi.

High-resolution clustering identified five microglial subclusters (MG1–MG5) with distinct distributions across experimental groups. The MG2 subset, which was prominently expanded in Q331K mice, was substantially reduced following cGAS inhibition (**Fig. 5b**). MG2 cells were enriched for *Apoe, Clec7a, Lyz2,* and *Fth1*, canonical markers of disease-associated microglia (**Fig. 5c**). Transcriptomic comparisons across total microglia revealed widespread reversal of disease-associated changes, with genes upregulated in Q331K-control mice relative to WT being largely downregulated by cGAS inhibition (**Fig. 5d-f**). This was reflected by a strong inverse correlation between cGASi response and Q331K-induced alterations (R = –0.9, p < 0.0001) (**Fig. 5g**).

Functionally, cGAS inhibition attenuated the hyperactivated microglial phenotype, suppressing DAM-associated genes (*Apoe, Trem2, Fth1*) while restoring the expression of homeostatic markers (*Cx3cr1, P2ry12*) (**Fig. 5h, Extended Data Fig. 4g**). Collectively, these findings indicate that cGAS inhibition reverses TDP-43 pathology-associated microglial overactivation and promotes a return toward homeostatic microglial states.

### cGAS inhibition restores oligodendrocyte maturation and myelin integrity in ALS mice

In addition to microglia, oligodendrocyte lineage cells and neurons also exhibited pronounced responses to TDP-43 pathology and cGAS inhibition. Disruption of oligodendrocyte lineage maturation and myelin integrity is a well-recognized feature of ALS pathology^38,39^. Given our observation that cGAS inhibition enhances microglial clearance of myelin debris, we next examined whether cGAS signaling also influences oligodendrocyte lineage dynamics in vivo. Using single-nucleus RNA sequencing data, we reconstructed oligodendrocyte developmental trajectories spanning from *Cspg4⁺* oligodendrocyte progenitor cells (OPCs), through *Enpp6⁺*/*Klk6⁺* immature oligodendrocytes (imOLs), to *Olig1⁺*/*Olig2⁺* mature oligodendrocytes (mOLs) (**Fig. 6a, b, Extended Data Fig. 5**). Q331K mice exhibited an accumulation of OPCs and imOLs accompanied by a reduction in mOLs, consistent with impaired oligodendrocyte lineage maturation, and this imbalance was partially normalized following cGAS inhibition (**Fig. 6c**).

**Figure 6.**
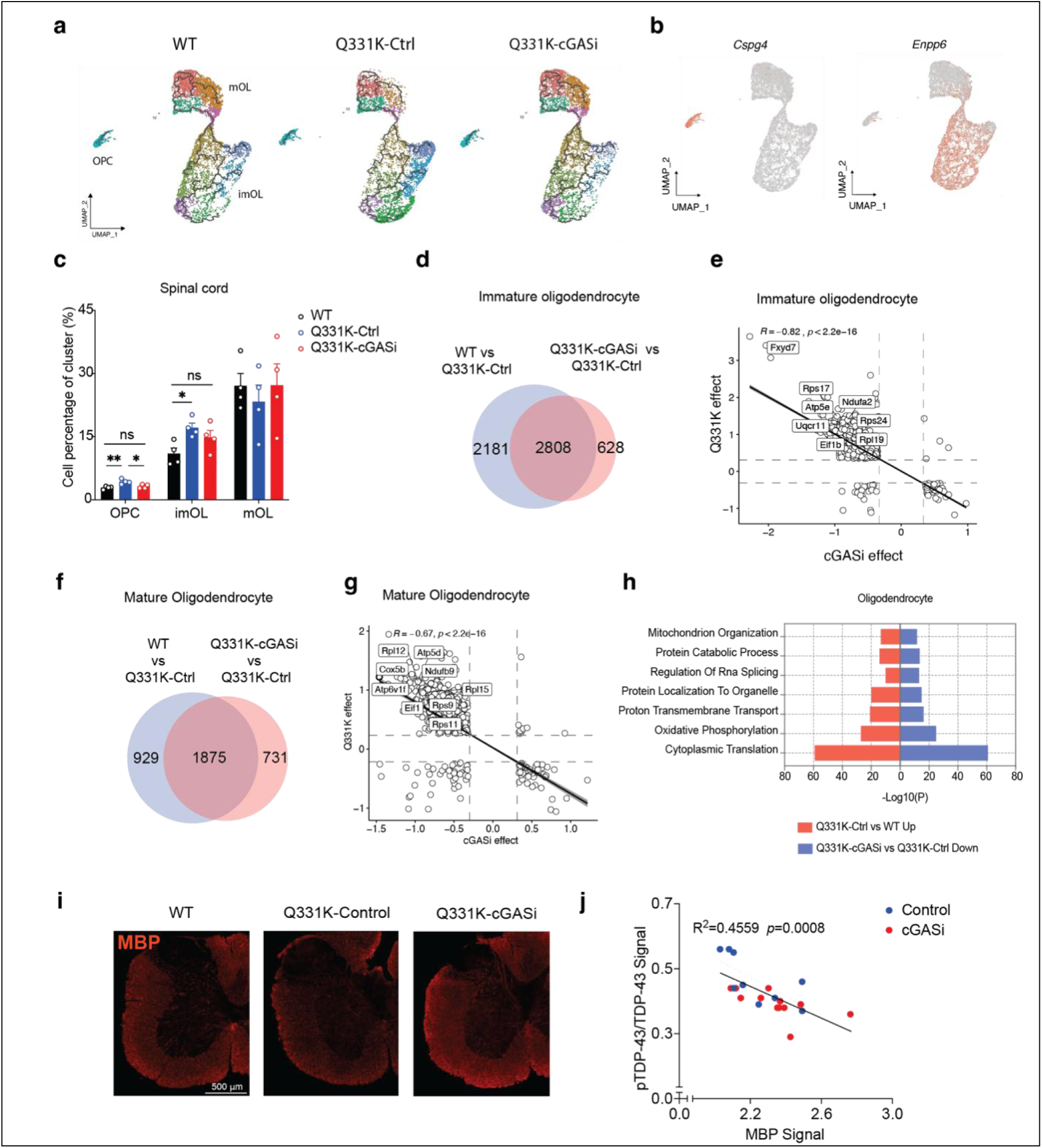
cGAS inhibition restores oligodendrocyte maturation in ALS mice. **a,** Pseudotime trajectory of oligodendrocyte-lineage cells (OPC, immature oligodendrocyte (imOL), and mature oligodendrocyte (mOL)) in the spinal cord of WT, Q331K-Ctrl, and Q331K-cGASi mice. **b,** UMAP plot of oligodendrocyte-lineage markers (*Cspg4* and *Enpp6*). **c,** The percentage of OPC, imOL, and mOL in the total cells of spinal cord in the single-nucleus analysis (WT, Q331K-Ctrl, Q331K-cGASi). N = 4 for WT-Control samples, n = 4 for Q331K-Control samples, and n = 4 for Q331K-cGASi samples. Data are presented as mean ± SEM. OPC Q331K-Control vs. WT-Control: \*\**P* = 0.0056; OPC Q331K-cGASi vs. Q331K-Control: \**P* = 0.0364; imOL Q331K-Control vs. WT-Control: \**P* = 0.0131. Data were analyzed by one-way ANOVA followed by a Tukey multiple comparison test; ns, not significant. **d,** Venn diagram for gene sets of imOL in the comparisons of WT vs. Q331K-Ctrl and Q331K-cGASi vs. Q331K-Ctrl. **e,** Correlation scatter plot with linear regression for DEGs of imOL under cGASi effect and Q331K effect. **f,** Venn diagram for gene sets of mOL in the comparisons of WT vs. Q331K-Ctrl and Q331K-cGASi vs. Q331K-Ctrl. **g,** Correlation scatter plot with linear regression for DEGs of mOL under cGASi effect and Q331K effect. **h,** Pathway enrichment bar plot for oligodendrocyte transcriptomes in the comparisons of Q331K-Ctrl vs. WT and Q331K-cGASi vs. Q331K-Ctrl. **i,** Representative images of MBP signal in the spinal cord of WT, Q331K-Ctrl, and Q331K-cGASi mice. **j,** Correlation plot relating MBP signal to pTDP-43/TDP-43 ratio in the spinal cord of Q331K mice with or without cGASi.

Differential expression analysis in imOLs revealed 2,808 overlapping genes between WT vs. Q331K-control and Q331K-cGASi vs. Q331K-control comparisons (**Fig. 6d**), with a strong inverse correlation between cGASi response and disease effect (R = –0.82, p < 0.0001) (**Fig. 6e**). A comparable transcriptional rescue was observed in mOLs, with 1,875 overlapping DEGs and a similarly strong inverse correlation (R = –0.67, p < 0.0001) (**Fig. 6f, g**).

Pathway enrichment analysis indicated that cGAS inhibition restored key oligodendrocyte-associated programs involved in ribosomal translation, oxidative phosphorylation, and RNA splicing (**Fig. 6h**), processes critical for myelin synthesis and maintenance. Consistently, immunofluorescence staining for myelin basic protein (MBP) revealed a significant negative correlation between MBP signal and spinal cord pTDP-43/TDP-43 ratio (R² = 0.456, p = 0.0008) (**Fig. 6i, j**), linking myelin deficits to the burden of TDP-43 pathology. Together, these data suggest that cGAS activation contributes to oligodendrocyte dysfunction in ALS, and that cGAS inhibition promotes oligodendrocyte maturation and myelin recovery in the degenerating spinal cord.

### cGAS inhibition restores RNA splicing homeostasis disrupted by TDP-43 pathology

TDP-43 dysfunction is defined by widespread disruption of RNA splicing, reflecting its canonical role as an RNA-binding protein and splicing regulator in neurons. We therefore asked whether cGAS signaling contributes to these core TDP-43–dependent splicing defects. Analysis of single-nucleus RNA sequencing data from mouse spinal cord neurons revealed a strong inverse relationship between Q331K-induced transcriptional changes and those elicited by cGAS inhibition, with splicing-related pathways among the most significantly affected (**Fig. 7a**).

**Figure 7.**
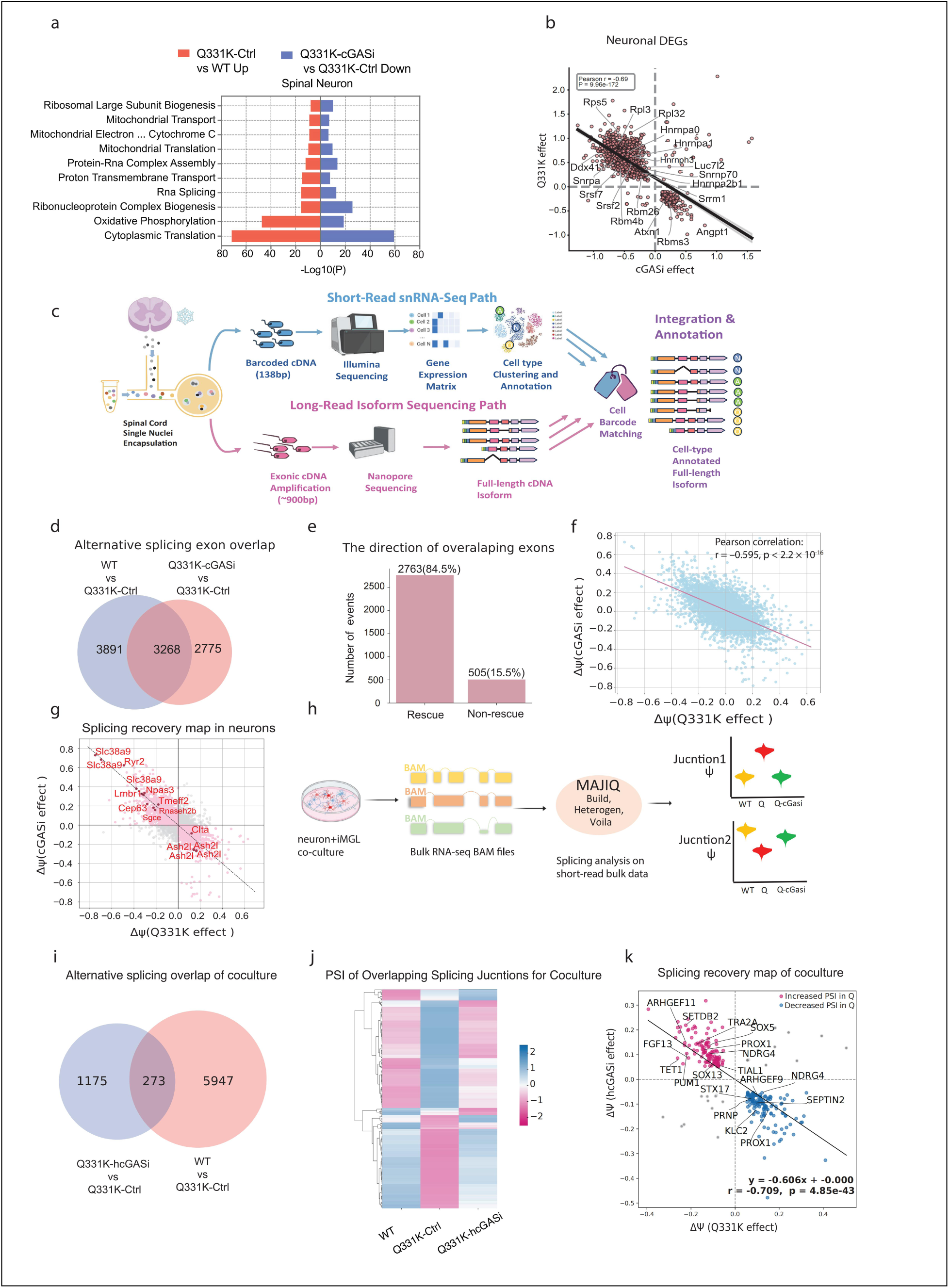
Inhibition of cGAS relieves TDP-43 pathology-induced RNA splicing defects in mouse and human ALS models. **a,** Pathway enrichment bar plot for neuronal transcriptomes in the comparisons of Q331K-Ctrl vs. WT (up) and Q331K-cGASi vs. Q331K-Ctrl (down) for spinal neurons. **b,** Scatter plot showing differential gene expression changes in spinal cord neurons identified by snRNA-seq. Each dot represents one differentially expressed gene (DEG), plotted by its effect size in TDP-43 Q331K mice (Q331K vs. WT, y-axis) and the effect size of cGAS inhibition (Q331K-cGASi vs. Q331K, x-axis). Notably, many genes showing opposing directional changes encode RNA-binding proteins. Selected representative genes are labeled. **c,** Illustration of SnISOr-Seq pipeline. The overall workflow is adapted in part from the SnISOr-Seq framework^40^, with modifications specific to this study. **d and e,** Alternative splicing events were aggregated across all annotated cell types to assess global overlap. In d, Venn diagram shows the overlap of alternatively spliced exons identified by SnISOr-Seq (|ΔΨ |>5%), indicating substantial overlap between disease-associated and cGAS inhibitor–responsive splicing events. In e, directionality analysis of the overlapping alternative exons showed 84.5% exhibited changes in opposite directions. Between the 2 comparisons, the observed bias toward opposite-direction changes was highly significant compared with random expectation (binomial test, p < 1 × 10⁻¹⁰). **f,** Scatter plot showing changes in ΔΨ for overlapping alternative splicing events identified by SnISOr-Seq across all cell types. Each point represents one alternative exon in a certain cell type that was detected in both comparisons. A strong negative correlation was observed between Q331K-induced and cGAS inhibitor–mediated changes (Pearson r = −0.595, p < 2.2 × 10⁻¹⁶). **g,** Scatter plot showing changes in ΔΨ for overlapping alternative splicing events identified by SnISOr-Seq in neurons. Selected genes are labeled. **h,** Illustration of MAJIQ pipeline. **i and j,** Alternative splicing events detected by MAJIQ in humanized co-culture system. In i, Venn diagram shows the overlap of alternative splicing events identified (|ΔΨ |>0.05, p<0.05 in t test). In j, the z-score heatmap showed the majority of the overlapping alternative junctions exhibited changes in opposite directions between the 2 comparisons. **k,** Scatter plot showing changes in ΔΨ for overlapping alternative splicing junctions identified by MAJIQ in co-cultured motor neurons (|ΔΨ |>5%, P<0.05 in t test). Each point represents one alternatively spliced junction detected in both comparisons. A strong negative correlation was observed between Q331K-induced and cGAS inhibitor–mediated changes (Pearson r = −0.709, p < 2.2 × 10⁻¹⁶).

Differential expression analysis further showed marked enrichment of RNA-binding protein genes among transcripts dysregulated by the TDP-43 Q331K mutation and selectively restored by cGAS inhibition (**Fig. 7b**). These findings prompted direct interrogation of RNA splicing regulation at isoform resolution.

To interrogate RNA splicing at single-exon resolution, we applied single-nucleus isoform RNA sequencing (SnISOr-Seq), which integrates long-read sequencing with single-nucleus transcriptomics to enable cell-type–resolved analysis of alternative splicing **(Fig. 7c**). Among captured exons across all cell types, we identified 3,268 alternative splicing events shared between disease-associated (Q331K-Control vs WT) and inhibitor-mediated (Q331K-cGASi vs. Q331K-Control) comparisons when applying a |ΔΨ| (delta PSI, percentage spliced in) threshold of 0.05. Strikingly, 84.5% of these exons exhibited reversed splicing direction following cGAS inhibition (**Fig. 7d, e**), with ΔΨ values showing strong negative correlation between Q331K and cGASi effects (R = –0.595, p < 0.0001) (**Fig. 7f**), demonstrating global rescue of TDP-43-driven splicing alterations.

Cell-type–resolved analysis revealed that rescued splicing events occurred predominantly in neurons and oligodendrocyte lineage cells (**Extended Data Fig. 6a, b**). In neurons, splicing recovery encompassed transcripts with diverse and essential functions (**Fig. 7g**). For example, *Ryr2*, encoding a ryanodine receptor critical for neuronal calcium homeostasis, and *Slc38a9*, a lysosomal amino acid transporter and nutrient sensor, which exhibited precise restoration to wild-type exon usage patterns following cGAS inhibition (*Ryr2*: WT Ψ = 0.83, Q331K-Control Ψ = 0.58, Q331K-cGASi Ψ = 0.82; *Slc38a9*: WT Ψ = 0.97, Q331K-Control Ψ = 0.60, Q331K-cGASi Ψ = 0.98) (**Extended Data Fig. 6c**).

To assess conservation of cGAS-dependent splicing regulation in human systems, we analyzed iPSC-derived motor neuron–microglial co-cultures using MAJIQ (Modeling Alternative Junction Inclusion Quantification) on bulk RNA sequencing data (**Fig. 7h, Extended Data Fig. 7**). We identified 273 alternatively spliced junctions shared between Q331K-induced and hcGASi-reversed events (**Fig. 7i**), with Z-score heatmap analysis revealing robust reversal of Q331K-driven splicing defects upon hcGASi treatment (**Fig. 7j**). Correlation analysis confirmed strong negative association between disease and inhibitor effects (R = –0.709, p < 0.0001) (**Fig. 7k**).

These findings demonstrate that cGAS inhibition restores TDP-43-associated RNA splicing homeostasis in a cell-type–specific manner across mouse and human model systems, targeting functionally essential neuronal transcripts.

## Discussion

In this study, we identify cGAS as a central link between TDP-43 pathology and glial and neuronal dysfunction in ALS. Using human postmortem tissue, TDP-43 mouse models, and human iPSC-derived systems, we show that cGAS signaling is consistently activated in TDP-43–associated disease states. cGAS activity drives pathological interactions between neurons and glia, contributing to microglial activation, lysosomal dysfunction, impaired oligodendrocyte maturation, neuronal RNA splicing abnormalities, and pathological TDP-43 phosphorylation. Notably, cGAS inhibition restores key aspects of RNA processing in both mouse and human TDP-43 models, alongside broader improvements in glial and neuronal function. These molecular and cellular effects translate into preserved motor performance and improved metabolic stability in ALS models.

Using both genetic ablation and pharmacological inhibition of cGAS in human iPSC-derived microglia, we uncovered a noncanonical role for cGAS in regulating lysosomal and phagocytic function that is independent of STING signaling. Previous studies have shown that STING activation promotes lysosomal biogenesis and autophagy through MiT/TFE family transcription factors, including TFEB and TFE3^41,42^. In contrast, our data show that inhibiting cGAS enhances microglial autophagic flux and lysosomal processing of myelin debris, accompanied by increased expression of the lysosomal protease cathepsin D (**Extended Data Fig. 2c, d**). These findings argue against a simple model in which cGAS acts solely upstream of STING to control lysosomal pathways. Instead, they are consistent with prior reports that cGAS directly interacts with LC3 via an LC3-interacting region to regulate micronucleophagy^43,44^. The lysosomal-promoting function of cGAS is consistent with the observation that cGAS inhibitor significantly ameliorated tau spread in a tauopathy mouse model^45^.

Remarkably, we showed that cGAS inhibitor rescued RNA splicing abnormalities that accompany TDP-43 pathology across multiple CNS cell types, with particularly prominent effects in neurons and oligodendrocyte lineage cells. Whether cGAS directly regulates RNA splicing or instead modulates splicing indirectly through inflammatory, transcriptional, or metabolic stress pathways remains unclear. Prior studies have shown that cGAS can localize to the nucleus and regulate DNA repair and chromatin-associated processes^24^, but the extent to which pharmacological cGAS inhibition affects these nuclear functions in neural cells is not known. Accordingly, while our findings raise the possibility that cGAS signaling may influence TDP-43–dependent RNA processing, delineating the precise molecular mechanisms will require targeted studies at the level of nuclear localization, chromatin state, and splicing factor engagement.

Our data support a model in which aberrant activation of the cGAS pathway contributes to TDP-43–driven ALS pathology through both cell autonomous and non-cell autonomous mechanisms. TDP-43–associated mitochondrial dysfunction promotes mitochondrial DNA release, providing a plausible upstream trigger for cGAS activation^16^. Our findings indicate that microglia serve as a major site of signal amplification. In vitro, transfer of neuronal cGAMP to microglia via volume regulated anion channels is sufficient to induce robust type I interferon responses^46^. Whether this intercellular signaling mechanism predominates in vivo, or operates in parallel with additional inflammatory or damage associated pathways, remains unresolved. Future studies will be required to determine whether cGAS inhibition can rescue TDP-43–associated splicing dysregulation in co-cultures of Q331K iMGLs and wild-type motor neurons.

Pharmacological inhibition of cGAS in human iPSC-derived microglia co-cultured with TDP-43 Q331K motor neurons partially restored neuronal RNA splicing fidelity, indicating that microglial innate immune activation can impair neuronal TDP-43 function in trans. This finding identifies a potential mechanism by which glial immune signaling exacerbates neuronal RNA dysregulation, a defining feature of ALS. Nevertheless, these co-culture systems represent a reductionist model and do not capture the full cellular complexity, temporal progression, or regional heterogeneity of the human spinal cord. Thus, the extent to which microglial cGAS activity drives neuronal splicing defects in vivo, particularly at later disease stages, warrants further investigation.

Beyond neurons and microglia, growing evidence indicates that oligodendrocytes are early and direct contributors to ALS pathogenesis. Oligodendrocyte-specific expression of mutant TDP-43 induces cell-autonomous dysfunction, myelin loss, and progressive motor impairment, particularly in the ventral spinal cord, accompanied by reduced expression of key myelin proteins^47,48^. Our finding that cGAS inhibition enhances oligodendrocyte maturation raises the possibility that cGAS signaling contributes to white matter pathology in ALS, either through intrinsic oligodendroglial mechanisms or indirectly via microglia-derived inflammatory cues.

Activated microglia impair oligodendrocyte survival and differentiation through the release of pro-inflammatory mediators, including TNF-α, IL-1β, and interferons, in part downstream of TLR7 signaling^49^. In parallel, microglia-derived reactive oxygen species exert disproportionate toxicity toward oligodendrocytes, which are particularly vulnerable due to their high metabolic demands and limited antioxidant capacity^50,51^. However, direct evidence for cGAS activation within oligodendrocytes remains limited, and cell-type-specific genetic approaches will be required to delineate the relative contributions of oligodendrocyte-autonomous versus microglia-mediated cGAS signaling.

In summary, we identify cGAS as a convergent signaling node linking TDP-43–associated cellular stress to multicellular dysfunction in ALS. In human iPSC-based systems, cGAS inhibition restores RNA splicing and normalizes neuron–glia interactions. In TDP-43 mouse models, cGAS inhibition improves RNA transcriptome and splicing programs, preserves motor behavior, and normalizes systemic metabolic dysfunction, but shows no efficacy in SOD1 ALS, indicating a mechanistically selective therapeutic effect. Together, these findings support development of brain-penetrant cGAS inhibitors for TDP-43–associated ALS, while underscoring the need to assess relevance to sporadic disease and long-term safety given cGAS’s role in antiviral immunity.

## Methods

### Cell culture

#### Human iPSC and THP-1 cell culture

Human iPSC lines carrying ALS-associated mutations were obtained from commercial sources: C9orf72 hexanucleotide repeat expansion (CS29iALS-C9nxx, Cedars-Sinai), C9orf72 isogenic control (CS29iALS-C9n1.ISOxx, Cedars-Sinai), TDP-43 (TARDBP) Q331K (JIPSC1064, The Jackson Laboratory). iPSC lines were maintained in mTeSR™ Plus medium (STEMCELL Technologies) on Matrigel-coated plates at 37°C with 5% CO₂.

THP-1 wildtype and cGAS knockout cell lines were maintained in RPMI-1640 medium (Thermo Fisher) supplemented with 10% heat-inactivated FBS (Gibco) and 1% penicillin-streptomycin (Life Technologies) at 37°C with 5% CO₂. Cells were passaged every 3-4 days to maintain logarithmic growth phase.

#### iPSC-derived iMGL differentiation, motor neuron differentiation and co-culture system

iMGLs were differentiated using STEMdiff™ Hematopoietic Kit and STEMdiff™ Microglia Differentiation & Maturation Kit (STEMCELL Technologies) following manufacturer protocols. iPSCs were induced toward mesoderm during the initial 3-day phase, followed by 9 days of hematopoietic differentiation with lineage-specific cytokines. Hematopoietic cells (HPCs) were harvested from culture supernatants and then transferred to Matrigel-coated plates and cultured in STEMdiff™ Microglia Differentiation Medium for 24 days. On day 24, cells were replated and matured in STEMdiff™ Microglia Maturation Medium for 4 days to generate fully mature iMGLs.

Motor neurons were differentiated using the STEMdiff™ Motor Neuron Kit (STEMCELL Technologies) following manufacturer protocols. On days 0-9, motor neuron progenitors were generated from iPSCs in AggreWell™ 400 plates. On days 9-14, motor neuron precursors underwent terminal differentiation to post-mitotic motor neurons. Then, the differentiated motor neurons were maintained in specialized motor neuron maturation medium until at least day 28 for full maturation.

For co-culture experiments, at day 28 of motor neuron differentiation, iMGLs were seeded at half the density of motor neuron precursors for day 9. Co-cultures were maintained in Microglia-Neuron Co-Culture Medium prepared by adding STEMdiff™ Microglia Supplement 2 to Motor Neuron Maturation Medium. Co-cultures were maintained for 4 days with half-medium changes every 2 days at 37°C with 5% CO₂.

#### Primary microglia culture

Primary microglial cells were isolated from postnatal day 1-3 mouse pups. Brain cortices were rapidly dissected, minced, and dissociated in 0.25% Trypsin-EDTA for 10 minutes at 37°C. DNase I was added to prevent cell clumping. Trypsin was neutralized with complete DMEM containing 10% heat-inactivated FBS, and cell suspensions were filtered through 70-μm cell strainers before centrifugation at 250g for 5 minutes. Mixed glial cultures were maintained in growth medium at 37°C with 5% CO₂ for 7-10 days. Following appearance of bright, round microglial cells, recombinant mouse GM-CSF (1 ng/mL, Life Technologies) was added to enhance microglial proliferation. After 48-72 hours, primary microglia were collected through mechanical agitation (210 rpm orbital shaker for 1 hour) and plated on poly-D-lysine-coated plates in DMEM supplemented with 10% heat-inactivated FBS and 1% penicillin-streptomycin. Experiments were performed 24 hours after plating to allow cellular adherence and recovery.

#### IncuCyte living imaging with myelin-pHrodo Red and LysoTracker

Quantitative assessment of microglial myelin uptake and lysosomal processing was performed using the IncuCyte system. At day −1, iMGLs were seeded at 4 × 10⁴ cells/well in a Matrigel-coated 96-well plate and incubate overnight at 37 °C with 5% CO₂. At day 0, cGAS inhibitor was added to a final concentration of 10 µM (with DMSO as control), followed by addition of LysoTracker Green (1 μM). 1 hour later, myelin debris conjugated to pHrodo Red (2 µg/ml) was added to each well. Plates were immediately transferred to the Incucyte system. Phase-contrast and fluorescence images were acquired at 1-hour intervals for 30 hours. In Incucyte analysis software, fixed background subtraction was applied, and fluorescence thresholds were set to quantify total integrated fluorescence intensity for each channel.

#### Animals

All animal procedures were approved by Institutional Animal Care and Use Committee of Weill Cornell Medicine. Mice were housed in groups of no more than five per cage, provided ad libitum access to food and water, and maintained in a pathogen-free barrier facility at 21–23 °C with 30–70% humidity on a 12-h light/12-h dark cycle. Transgenic mouse lines were obtained from the Jackson Laboratory: Q331K (B6.Cg-Tg(Prnp-TARDBP*Q331K)103Dwc/J, JAX stock #017933), A315T (B6;C3-Tg(Prnp-TARDBPA315T)95Balo/J, JAX stock #010700), and SOD1 (B6SJL-Tg(SOD1G93A)1Gur/J, JAX stock #002726).

Mouse cGAS inhibitor (mcGASi, TDI-6570) was formulated into standard rodent chow at 150 mg/kg concentration by Research Diets Inc using the methods as described previously ^15^. Control diet pellets were prepared identically without drug incorporation. Drug treatment was initiated at 6 weeks of age and continued throughout the experimental period. All diet pellets were stored at 4°C and used within 6 months of preparation to ensure compound stability.

#### Synthesis of cGAS inhibitors

TDI-6570 and SS-1386 were prepared using the methods as described previously^34^. All intermediates were obtained by filtration using water and cold methanol. Purity of the products was confirmed by performing liquid chromatography–mass spectrometry analysis.

#### Cell-free cGAS activity assay

Activities of compounds SS-1386 were determined by measuring the conversion of ATP and GTP to cGAMP in the presence of dsDNA in a reaction buffer composed of Tris-HCl (20 mM, pH 7.4), NaCl (150 mM), MnCl2 (0.2 mM, human cGAS) and Tween 20 (0.01%) and the remaining ATP concentration using a Kinase-GloMax luminescent kinase assay (Promega), as previously described^15^. Briefly, 10 μl of a master mix of 0.4 mM ATP, 0.4 mM GTP and 0.02 mg/ml dsDNA in the reaction buffer supplemented with 2 mM DTT was added to reaction wells containing SS-1386 in the same buffer (20 μl). Next, 10 μl of a 4Å∼ human cGAS (0.4 μM) solution in the reaction buffer supplemented with 2 mM DTT was added to appropriate wells. Similar sets of reactions without cGAS or the inhibitor were set by adding the buffer alone. The reactions with the plates sealed were incubated at 37 ÅãC and stopped by the addition of 40 μl of

Kinase-GloMax. Luminescence was recorded in relative light units (RLUs) using a Biotek Synergy H1 hybrid plate reader (BioTek). ATP depletion was normalized against the positive control (no cGAS) and negative control (with cGAS) as follows: percent inhibition = 100 Å∼ (RLU sample − RLU average negative control) ÅÄ (RLU average positive control – RLU average negative control).

#### Western blot analysis

Cells and tissues were rapidly lysed in ice-cold RIPA buffer supplemented with protease inhibitor cocktail and phosphatase inhibitor cocktails (Millipore Sigma). Samples were incubated on ice for 30 minutes, and then centrifuged at 14,000g for 10 minutes at 4°C. Supernatants were collected, and protein concentrations were determined using Pierce BCA Protein Assay Kit (Thermo Fisher). Equal amounts of protein samples were separated on SDS-PAGE gels, and transferred to nitrocellulose membranes. Membranes were blocked with 5% non-fat milk in TBS with 0.01% Triton X-100 for 1 h at room temperature, and then incubated overnight at 4°C with primary antibodies. Following three 10-minute washed in TBST, membranes were incubated with HRP-conjugated secondary antibodies for 1 hour at room temperature. Bands were visualized using enhanced chemiluminescence (Bio-Rad) and were quantified using ImageLab (Bio-Rad) and FIJI (NIH) software.

#### Immunofluorescence

iPSCs and iPSC-derived cells were fixed in 4% paraformaldehyde diluted in PBS for 15 minutes at room temperature and then washed three times with DPBS. Cells were permeabilized in 0.1% Triton X-100 in DPBS for 10 minutes, followed by blocking in solution containing 0.1% Triton X-100 and 5% bovine serum albumin in DPBS for 1 hour at room temperature. Primary antibodies diluted in blocking solution were applied overnight at 4°C. Following three 5-minute washes with DPBS, secondary antibodies conjugated with Alexa Fluor fluorophores (1:500, Invitrogen) were applied for 1 hour at room temperature in the dark. Coverslips were washed three times with DPBS and mounted on slides. For tissue immunofluorescence, spinal cords were fixed in 4% paraformaldehyde for 48 h, and them cryoprotected in 30% sucrose in PBS for 48h at 4 °C until tissues sank. Spinal cord segments were embedded in OCT compound (Sakura), cut using a freezing microtome (Leica), and placed in cryoprotective medium at –20 °C until use.

Tissue sections were processed using similar staining protocols as described for cultured cells. Images were acquired using Apotome (Zeiss), Keyence (BZ-X710), and LSM 880 Laser Scanning Confocal Microscope (Zeiss). Image processing and quantification were performed using Zen 3.2 software (Zeiss) and FIJI (NIH).

#### Antibodies

Primary antibodies used included: anti-cGAS (1:500, Cell Signaling #15102), anti-phospho-TBK1 Ser172 (1:500, Cell Signaling #5483), anti-TBK1 (1:1000, Cell Signaling #3504), anti-phospho-TDP-43 (1:1000, ProteinTech #22309-1-AP), anti-TDP-43 (1:1000, ProteinTech #10782-2-AP), anti-APOE (1:1000, Millipore #AB947), anti-CTSD (1:1000, Abcam #ab75852), anti-CHAT (1:500, Millipore #AB144P), anti-TMEM119 (1:200, Abcam #ab209064), anti-IBA1 (1:500, WAKO #019-19741 or Abcam #ab5076), anti-MBP (1:1000, Millipore #AB5864), anti-β-actin (1:5000, Sigma-Aldrich #3854), and anti-GAPDH (1:5000, GeneTex #GTX100118).

#### Bulk RNA sequencing

Total RNA was isolated from cells using RNeasy Mini Kit (Qiagen) according to manufacturer protocols. RNA concentration was measured using NanoDrop. Library preparation and bulk RNA sequencing were performed by Novogene by sequencing on NovaSeq platform. Gene-level counts were quantified using featureCounts, and differential gene expression analysis was performed in R using DESeq2 package. Visualization and pathway analysis were performed using ggplot2 package, and pathway analysis was performed using Gene Set Enrichment Analysis software.

#### Single-cell and single-nucleus RNA sequencing and analysis

Single-cell RNA sequencing (scRNA-seq) data of isolated live human microglia from ALS patients were obtained from a previously published study by the Olah lab^33^. Raw count matrices and associated metadata were downloaded from GEO GSE204704. We also performed single-nucleus RNA-sequencing (snRNA-seq) on spinal cords from wild type (WT), Q331K and Q331K-cGASi transgenic mice. Briefly, spinal cord tissues were rapidly dissected, flash-frozen in liquid nitrogen, and stored at −80°C until processing. Nuclei were isolated using modifications of established protocols^15,49^. Libraries were prepared using Chromium Single Cell 3’ Reagent Kits (10x Genomics) and sequenced on NovaSeq 6000 (Illumina). Raw sequencing data were processed using Cell Ranger software to align reads and generate gene-barcode matrices. Quality control removed genes expressed in fewer than two cells, nuclei with unique gene counts below 200 or above 4,000, and nuclei with mitochondrial read fractions exceeding 5%. Normalization, dimensionality reduction, and clustering were performed using Seurat package in R. Principal component analysis was performed on all genes, with t-distributed stochastic neighbor embedding was run on the top 20 principal components. Cell clusters were identified using Seurat FindNeighbors and FindClusters functions. Differential gene expression analysis employed FindMarkers function. For the Q331K cohort snRNA-seq analysis, we integrated and analyzed samples from WT mice (n=4), Q331K-Control mice (n=4), and Q331K-cGASi mice (n=4).

### Alternative splicing analysis

#### Short-read splicing analysis (MAJIQ)

Alternative splicing analysis of short-read bulk RNA sequencing data was performed using MAJIQ (v2.5.8)^52^. STAR-aligned BAM files generated from sequencing reads against GRCh38 primary human genome with GENCODE v44 annotations as reference index were used as input for MAJIQ build module. Using the GENCODE v44 GFF3 annotation, MAJIQ build produced a splicing graph and individual junction files for each sample (3 WT, 3 Q331K and 3 cGASi) based on the GRCh38 reference genome. Changes in local splicing variations between conditions were quantified using the MAJIQ Heterogen module, which accounts for variability across samples rather than assuming uniform splicing patterns within each condition. MAJIQ Heterogen outputs were exported as tab-separated value (TSV) files using the MAJIQ tsv module, with local splicing events annotated. These files were then processed using custom Python scripts to extract Ψ values and the associated probabilities of splicing change for each junction. Junctions were considered differentially spliced if they met both a statistical significance threshold (t-test, p < 0.05) and an absolute change in Ψ greater than 5% between conditions. Overlapping junctions were defined as junctions mapping to the same gene with identical genomic coordinates that were detected in both comparisons (WT vs Q331K and Q331K vs hcGASi).

#### Long-read splicing analysis (SnISOr-Seq)

##### Library prep

Splicing analysis for mouse spinal cord samples was performed through SnisorSeq^40^. The SnISOr-Seq library preparation was performed as previously described^53^, with minor modifications. Briefly, an exome capture step was applied to enrich for spliced cDNA molecules using the SureSelectXT Mouse All Exon Probe (Agilent, 5190-4641). Library preparation was otherwise carried out according to established SnISOr-Seq protocols.

##### Long-read alignment

Long-read sequencing data were processed as previously described^53^. Reads were aligned to the mouse reference genome (GRCm39) using Minimap2 (v2.24).

##### Differential exon usage analysis and Ψ calculation

To correct for PCR-induced molecular duplication, reads containing unique molecular identifiers (UMIs) within an edit distance of 4 from more abundant UMIs were discarded. Alternative exons were identified from the remaining long reads.

For each exon passing quality filters, exon inclusion and exclusion events were quantified separately for each cell type and experimental condition (scisorATAC package^54^). These values were summarized in 2 × 2 contingency matrices to evaluate differential exon usage. Statistical significance was determined using Fisher’s exact test, and no significance testing was performed when contingency tables did not satisfy the chi-squared applicability criterion. To control for multiple comparisons, false discovery rates were estimated using the Benjamini–Yekutieli procedure. Ψ values and ΔΨ were derived from the inclusion and exclusion counts. Only exons with Ψ values between 1% and 90% and supported by at least 2 reads per condition were retained for downstream differential splicing analyses.

##### Visualizing long reads

Long-read data was visualized using ScisorWiz (version 1.2.1.2)^55^.

#### NFL ELISA for serum

NFL levels were measured in serum samples using Neurofilament Light Chain ELISA Kit (Quanterix) according to manufacturer protocols. Serum samples were collected via cardiac puncture, allowed to clot at room temperature for 30 minutes, and centrifuged at 1,000g for 10 minutes at 4°C. Serum was separated and stored at −80°C until analysis. Samples were diluted in assay buffer and analyzed in duplicate according to kit instructions.

#### Behavioral and metabolic tests

##### Rotarod

Motor coordination was assessed using accelerating rotarod apparatus (Ugo Basile) with programmed acceleration from 4 to 40 rpm in 5 minutes. Mice were habituated to the testing room for 1 hour prior to assessment. During initial training phase, mice were habituated to the apparatus for three consecutive days with two trials per day. Formal testing commenced on the fourth day and was repeated every 2 weeks throughout the experimental period. Latency to fall was recorded manually when automatic sensors failed to detect lightweight mice.

##### Metabolic cage

Metabolic parameters were assessed using Promethion metabolic cage system. Mice underwent single-housing adaptation for 3-4 days before data collection to minimize stress responses associated with social isolation. Following adaptation, mice were transferred to metabolic chambers for continuous monitoring. The system continuously recorded oxygen consumption, CO₂ production, energy expenditure, food intake, water intake, locomotor activity, and body position. Energy expenditure was derived from indirect calorimetry using the Weir equation and normalized to lean body mass quantified via EchoMRI. Circadian analysis segregated data into 12-hour light and dark phases. All environmental parameters were maintained constant throughout monitoring periods.

##### Hindlimb

Mice were held by the tail and lifted with its hindlimbs observed for retraction towards the abdomen. Motor function decline was assessed using a 5-point neurological scoring system: 0 = normal motor function; 1 = tremor and/or abnormal hind limb posture; 2 = moderate gait abnormalities; 3 = severe gait abnormalities with dragging of hind limbs; 4 = inability to right when placed on back; 5 = complete hind limb paralysis. Assessments were performed weekly by blinded observers with inter-rater reliability validation.

## Data availability

All data associated with this study are included in the paper and the Supplementary Information. RNA-seq data have been deposited in the Gene Expression Omnibus (GEO) under BioProject accession number PRJNA1403051. All the data will be deposited in GEO and made publicly available prior to publication.

## Code availability

All custom code used for sequencing analysis are publicly available at https://github.com/lifan36. Any additional analysis scripts required to reanalyze the data reported in this study is available from the corresponding author upon reasonable request.

## Acknowledgments

We thank Rohit Sharma and Tara Doma Lama at the Metabolic Phenotyping Center of Weill Cornell Medicine for conducting the metabolic cage experiments. The study was supported by NIH R01AG072758 (to L.G.), R01AG074541 (to S.C.S. and L.G.), 1R01AG079291-01A1 (to L.G.), R01AG079557-01 (to L.G.), and Freedom Together Foundation (to L.G.); Rainwater Charitable Foundation (to L.G.); Cure Alzheimer Fund (to S.C.S. and L.G.).

## Author Contributions

Conceptualization: L.G. and Y.L.; investigation, Y.L., W.F., A.A., S.-I.L., L.F., M.B., R.K.N., M.Y.W., S.A., J.Z., S.W, P.Y., and K.N.; analysis, Y.L., A.A., W.F., and L.F.; funding acquisition, L.G., and S.C.S.; resources, M.O., L.F., and H.T.; visualization, Y.L., W.F., A.A., L.F., R.K.N., W.Q., and L.G.; writing—original draft, Y.L., W.F., A.A., and L.G..

## Notes

### Competing Interest Statement

The authors have declared no competing interest.

