## Supplementary_figures for "cGAS inhibition delays TDP-43-driven ALS Pathogenesis"

### Supplementary Materials

### Extended Figures 1–7

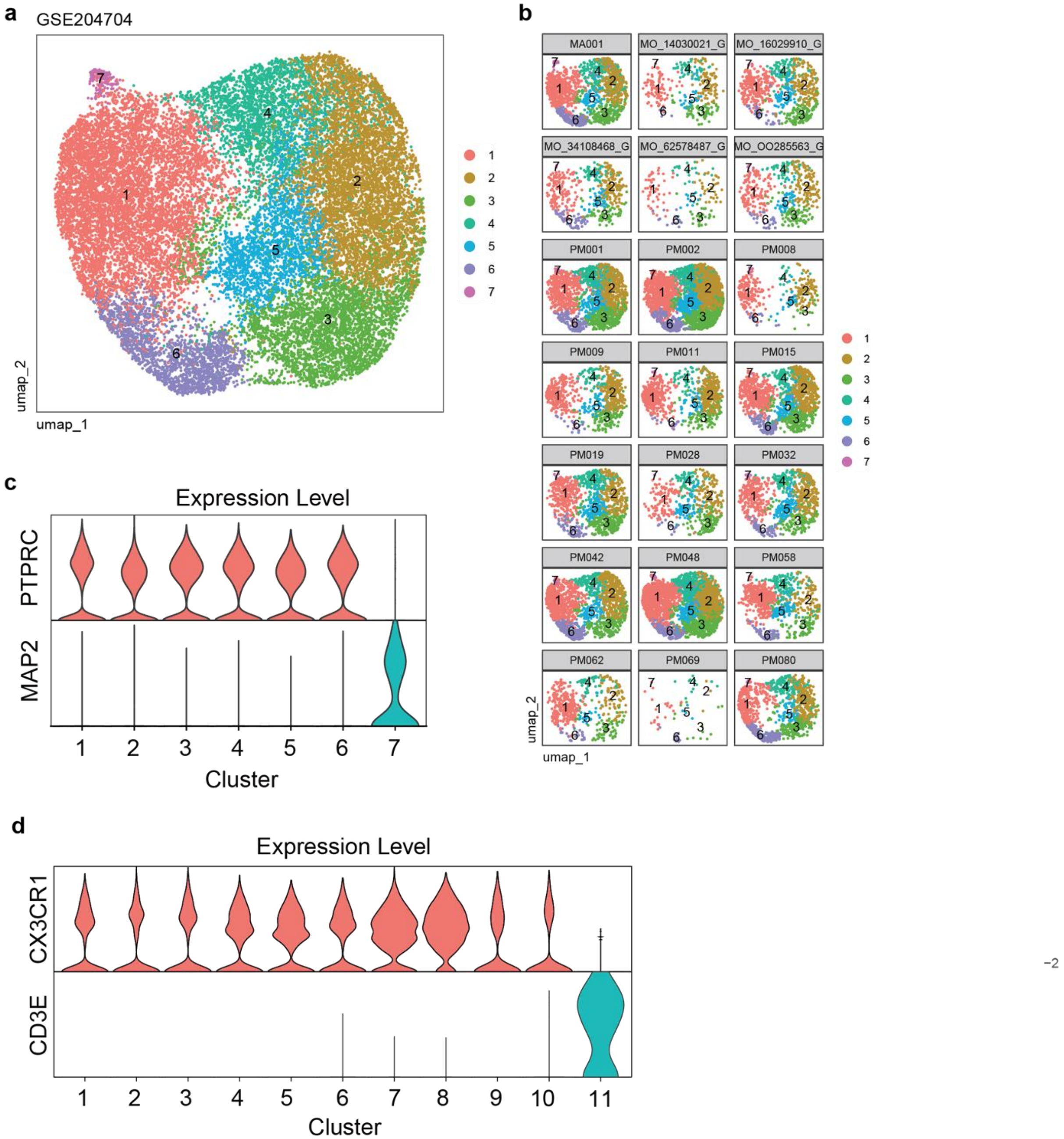


**Extended Data Fig. 1 Single cell sequencing analysis of human ALS and non-ALS brains. a,** UMAP plot of cells from the ALS and non-ALS brain samples. **b,** UMAP plot of each human brain samples. **c,** Violin plot of *PTPRC* (CD45) and *MAP2* expression in cells from cluster 1-7. **d,** Violin plot of *CX3CR1* and *CD3E* expression in immune cells from subcluster 1-11.

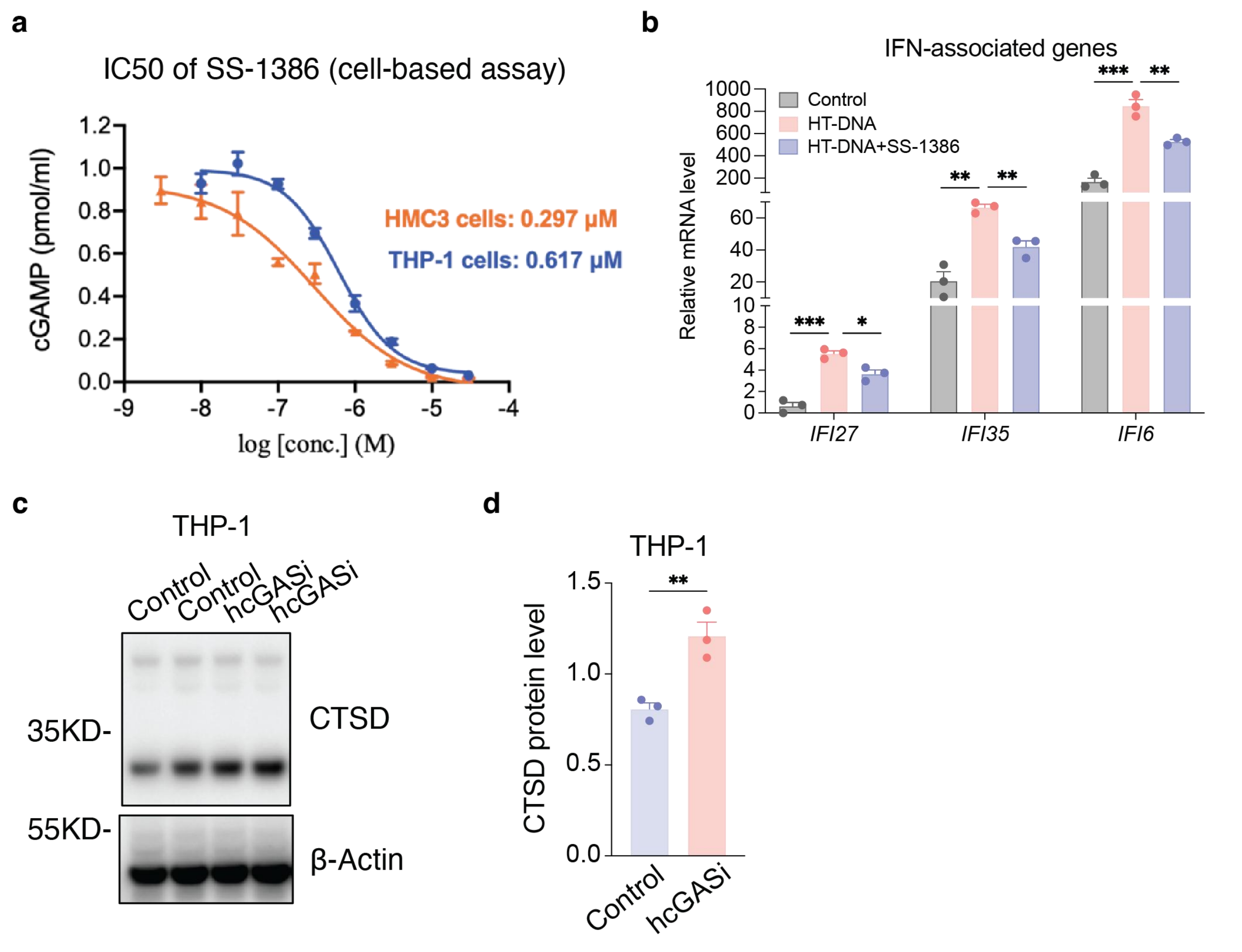


**Extended Data Fig. 2 Activity and specificity of the human cGAS inhibitor SS-1386. a,** Dose-dependent inhibition of HT-DNA induced cGAMP production in THP-1 dual reporter line and HMC3 microglial line. THP1: IC50 = 0.62 µM and HMC3: IC50 = 0.30 µM. Data are reported as mean ± SEM. n = 2 per condition. **b,** Bar plots of mRNA levels of interferon-associated genes in iMGLs (Control, HT-DNA and HT-DNA- SS-1386). N = 3 for Control samples, n = 3 for HT-DNA samples, and n = 3 for HT-DNA+SS-1386 samples. Data are presented as mean ± SEM. *IFI27* HT-DNA vs. Control: ****P* = 0.0001; *IFI27* HT-DNA+SS-1386 vs. HT-DNA: **P* = 0.0138; *IFI35* HT-DNA vs. Control: ***P* = 0.0017; *IFI35* HT-DNA+SS-1386 vs. HT-DNA: ***P* = 0.0038; *IFI6* HT-DNA vs. Control: ****P* = 0.0001; *IFI6* HT-DNA+SS-1386 vs. HT-DNA: ***P* = 0.0057. Data were analyzed by two-way ANOVA with a Tukey multiple comparisons test. **c and d,** Representative immunoblots (c) and quantification (d) of CTSD and cGAS antibodies in THP-1 cells with or without hcGASi , normalized to β-Actin. N = 3 for Control samples, and n = 3 for hcGASi samples. Data are presented as mean ± SEM. hcGASi vs. Control: ***P* = 0.0085. Data were analyzed by two-tailed unpaired t-test.

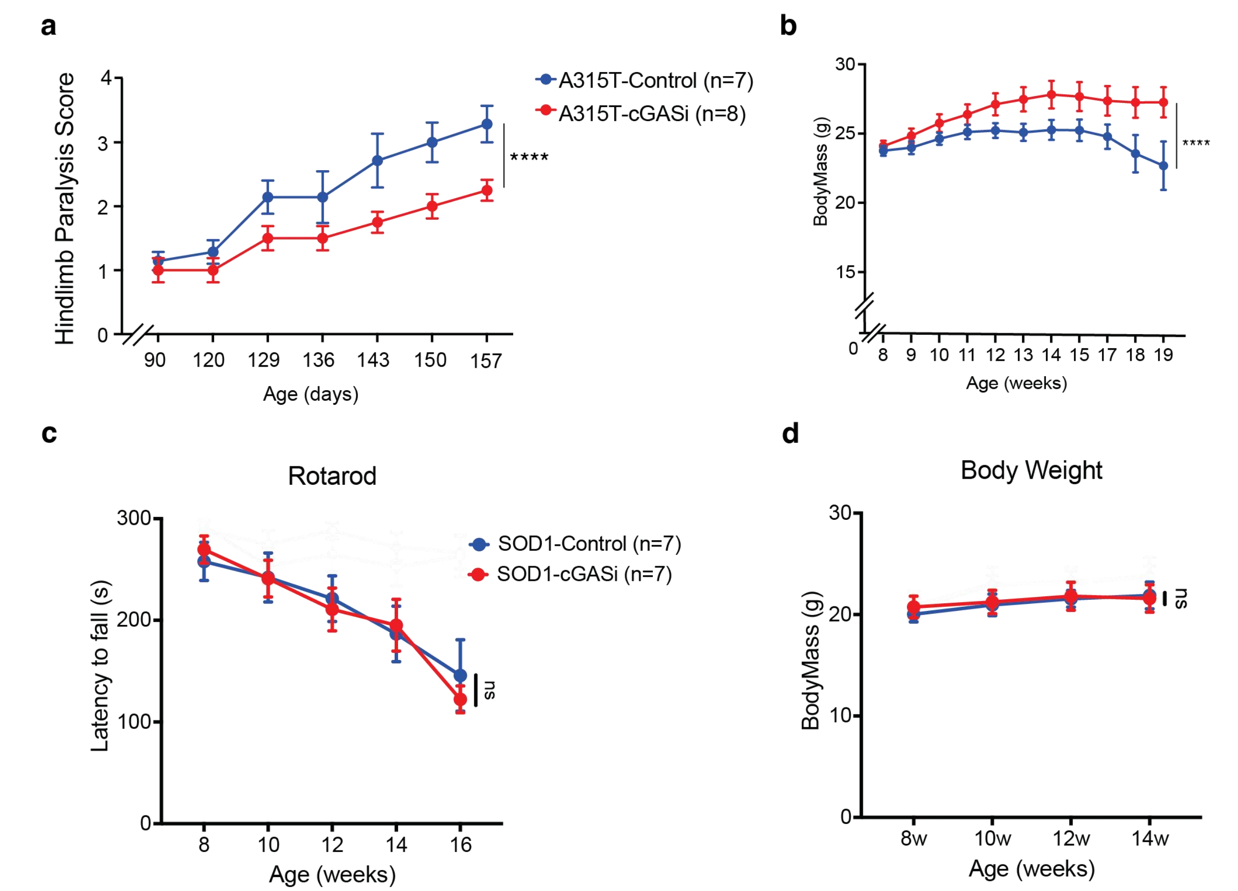


**Extended Data Fig. 3 Behavioral tests for A315T and SOD1 mice. a,** Hindlimb paralysis score over time in A315T mice with or without cGASi. N = 7 for A315T-Control samples, and n = 8 for A315T-cGASi samples. Data are presented as mean ± SEM. A315T-cGASi vs. A315T-Control: *****P* < 0.0001. Data were analyzed by two-way ANOVA with a Tukey multiple comparisons test. **b,** BodyMass (g) over time in A315T mice with or without cGASi. N = 7 for A315T-Control samples, and n = 8 for A315T-cGASi samples. Data are presented as mean ± SEM. A315T-cGASi vs. A315T-Control: *****P* < 0.0001. Data were analyzed by two-way ANOVA with a Tukey multiple comparisons test. **c,** Rotarod latency to fall (s) across ages from 8 weeks to 16 weeks for SOD1 mice with or without cGASi. N = 7 for SOD1-Control samples, and n =7 for SOD1-cGASi samples. Data are presented as mean ± SEM. Data were analyzed by two-way ANOVA with a Tukey multiple comparisons test; ns, not significant. **d,** BodyMass (g) over time in SOD1 mice with or without cGASi. N = 7 for SOD1-Control samples, and n =7 for SOD1-cGASi samples. Data are presented as mean ± SEM. Data were analyzed by two-way ANOVA with a Tukey multiple comparisons test; ns, not significant.

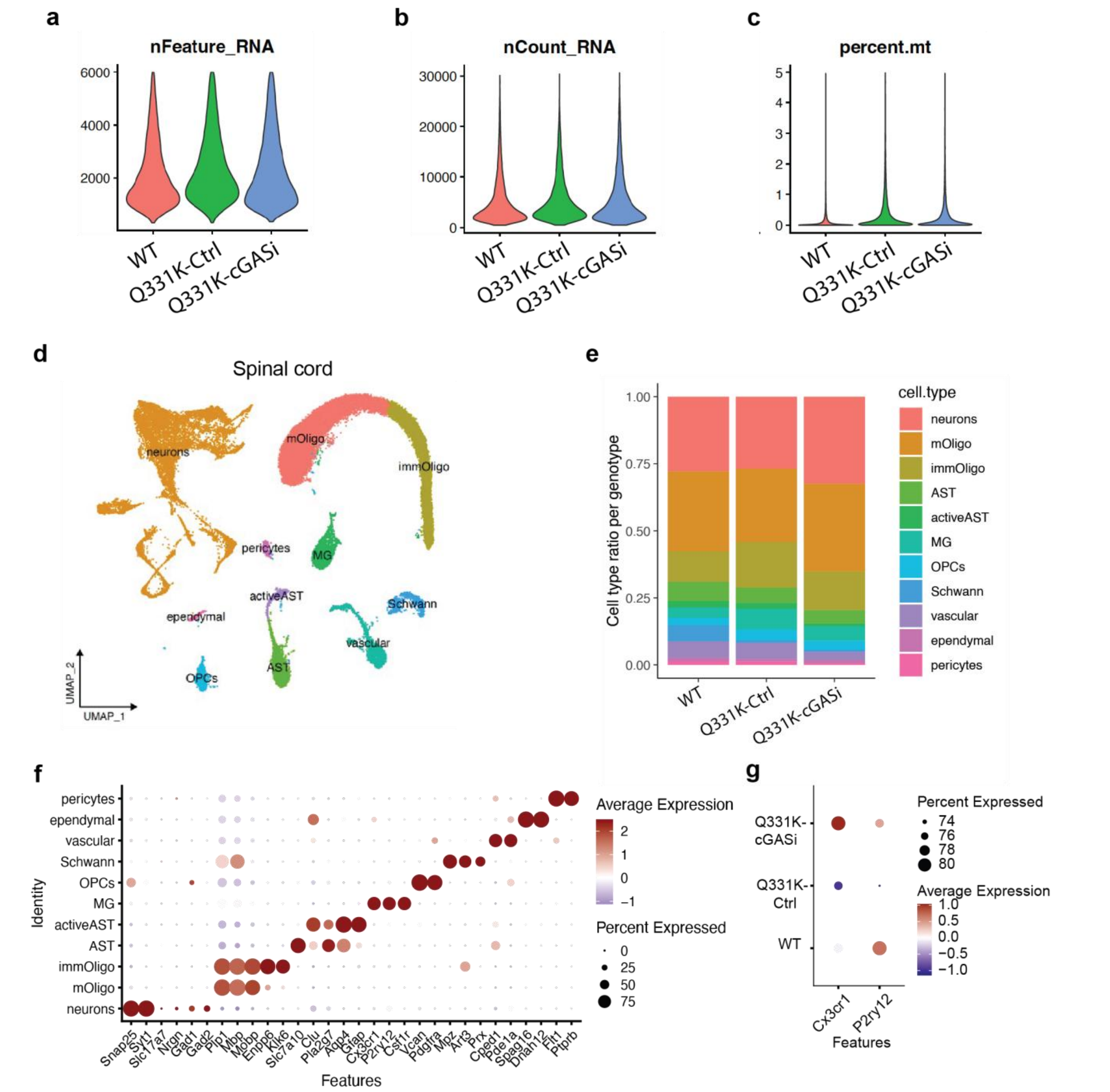


**Extended Data Fig. 4 Single nucleus sequencing analysis for spinal cord of Q331K mice. a–c,** Quality control feature plots from single nucleus RNA-seq of spinal cord tissues: a, number of detected genes per nucleus (nFeature_RNA); b, total UMI (unique molecular identifier) counts per nucleus (nCount_RNA); c, percentage of mitochondrial transcripts per nucleus (percent.mt). **d,** UMAP plot for cells of spinal cord tissues. **e,** Bar plot for cell type ratios in spinal cord tissues of WT, Q331K-Ctrl, Q331K-cGASi groups. **f,** Dotplot for gene markers of central cell types in spinal cord tissues. **g,** Dotplot for *Cx3cr1* and *P2ry12* in spinal microglia in WT, Q331K-Ctrl, and Q331K-cGASi.

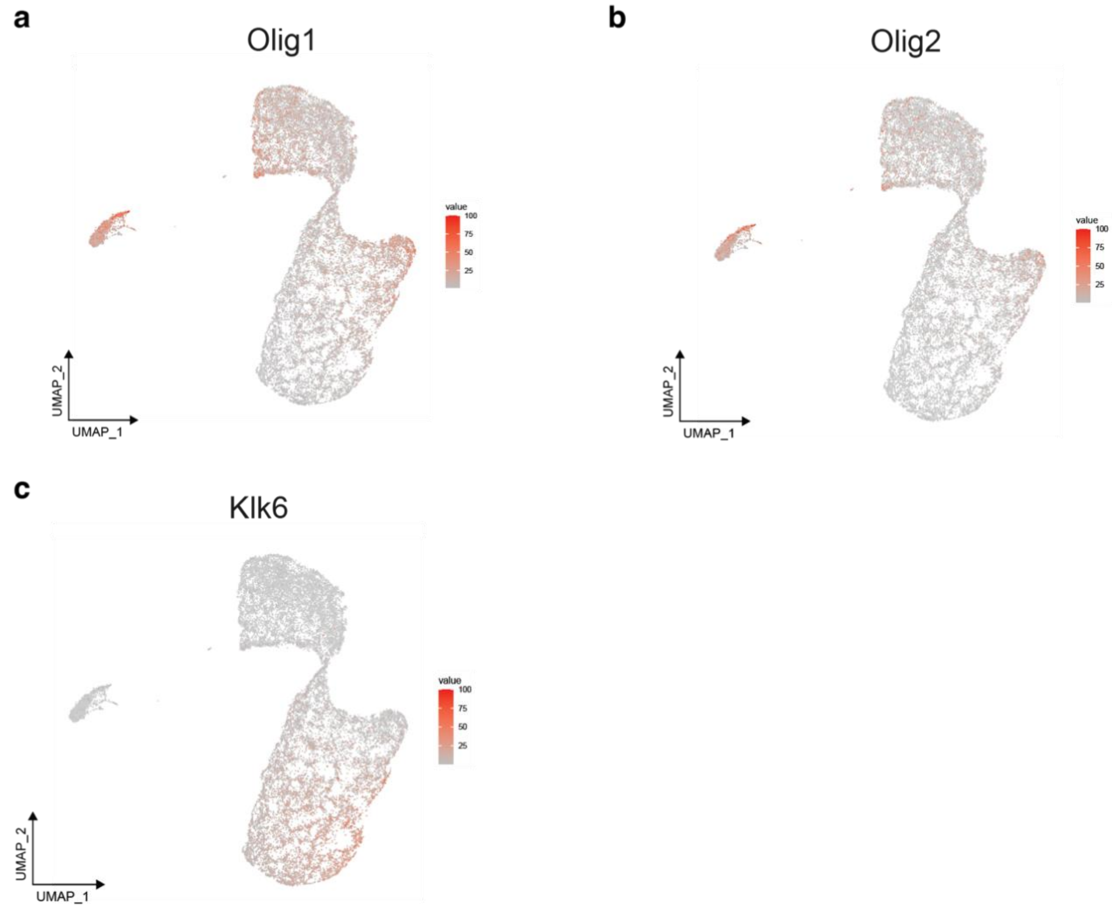


**Extended Data Fig. 5 Single nucleus sequencing analysis for oligolineage in Q331K mice.**
**a–c,** UMAP plots of oligodendrocyte-lineage markers: a, *Olig1*; b, *Olig2*; c, *Klk6*.

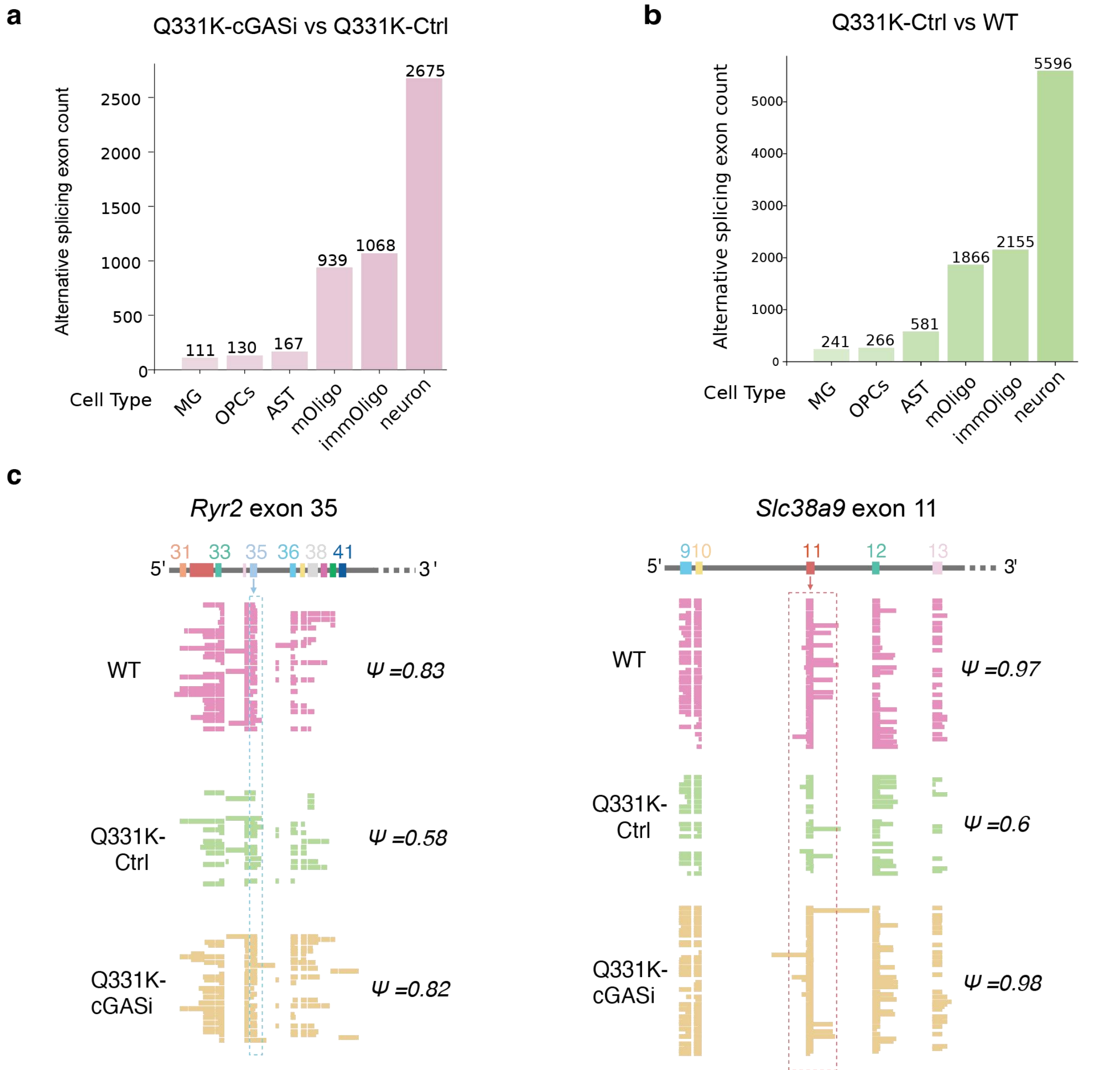


**Extended Data Fig. 6 RNA splicing analysis for spinal cord of Q331K mice. a and b,** Number of alternatively splied exons identified by SnISOr-Seq across major spinal cord cell types in Q331K mice treated with cGAS inhibitor compared with untreated Q331K controls and Q331K mice compared with wild-type controls (|ΔΨ |>0.05). Bars indicate the total number of alternative exons detected in each cell type. **c,** Visualization by SciSOrWiz to show isoform changes across conditions for representative genes.

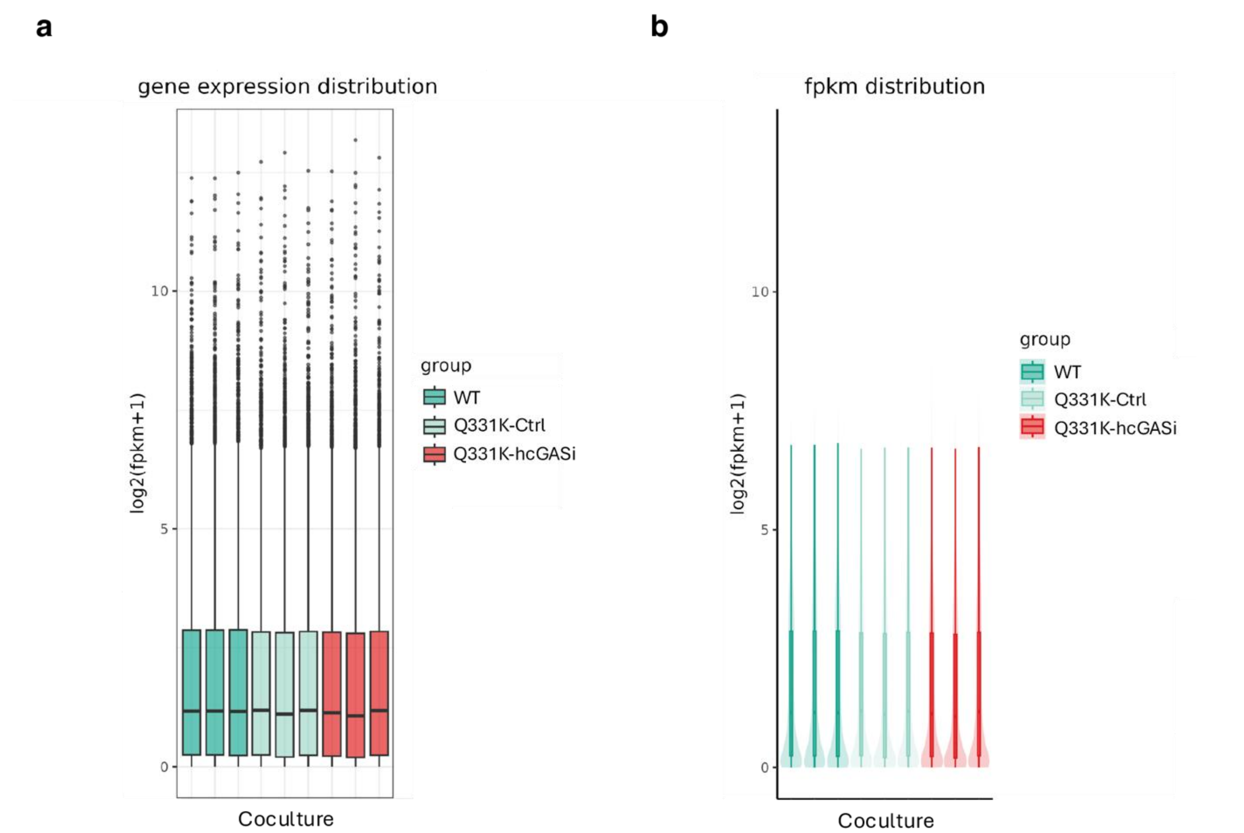


**Extended Data Fig. 7 Bulk RNA-seq analysis of iPSC-derived co-culture system. a,** Bar plot for gene expression distribution of bulk RNA-seq samples across WT, Q331K-Ctrl, and Q331K-cGASi. **b,** Bar plot for fpkm distribution of bulk RNA-seq samples across WT, Q331K-Ctrl, and Q331K-cGASi.
